# Cellular coding of ingestion in the caudal brainstem

**DOI:** 10.64898/2025.11.30.691333

**Authors:** Truong Ly, Xinyang Yi, Grace R. Lee, James CR Grove, Youssef E. Sibih, Jun Y. Oh, Longhui Qiu, Ho Namkung, Nilla Sivakumar, Zachary A. Knight

## Abstract

The termination of a meal is triggered by sensory feedback from the stomach and intestines that reports on ingested food^1–10^ and is relayed to the caudal nucleus of the solitary tract (cNTS) in the brainstem^11–14^. This sensory feedback is thought to gradually intensify as a meal progresses, resulting in the progressive activation of cNTS circuits that promote satiety^15,16^, but this idea has never been tested by recording the single-cell activity of cNTS neurons while animals eat. Here, we have used a preparation for calcium imaging in the caudal brainstem of behaving animals^17^ to characterize how food ingestion is encoded in the cNTS. We find that when food is delivered directly to the stomach, most cNTS neurons exhibit ramping activation over many minutes that tracks cumulative food consumed and depends on canonical gut-brain pathways. However, when the same food is consumed by mouth, this widespread ramping activation is gone, and most cNTS neurons instead exhibit phasic, seconds-timescale responses to oral contact with food. We show that these rapid responses are driven by a combination of mechanical, gustatory and nutritive signals from the mouth and throat and do not require traditional gut-brain pathways, including gut-innervating vagal afferents, although GI feedback modulates their duration. We show that one source of this rapid input is descending projections from the paraventricular hypothalamus, which track ingestion dynamics and are required for proper meal termination. These findings reveal that sensory feedback from the stomach and intestines, which is directly transmitted to the cNTS and thought to be critical for satiety, is not the major driver of cNTS activity during ingestion. Instead, these circuits make extensive use of rapid, pregastric signals that report on the dynamics of behavior.

## Introduction

The caudal nucleus of the solitary tract (cNTS) is the direct target of the vagal and glossopharyngeal sensory neurons that innervate the entire alimentary canal from the oropharynx to the intestines^5,6,18–20^. These sensory neurons track the volume and nutrient content of ingested food and then relay this information to the cNTS, a brainstem structure that is critical for meal termination^21–29^. The cNTS also senses circulating hormones and nutrients, both directly and indirectly through the adjacent area postrema^30,31^. These neural and blood-borne signals of food ingestion are thought to gradually intensify as a meal progresses, in a manner that tracks cumulative food consumed, thereby leading to the progressive activation of the cNTS circuits that promote satiety^15,16^. Thus, the cNTS is the first station in the brain that receives many visceral signals of food consumption, and it uses these signals to determine when an animal should stop eating^1–3,17,32^.

Because the cNTS is a primary entry point for visceral feedback in the brain, the regulation of its neurons during eating provides a window into how the sensory signals generated during a meal are used to control behavior. However, little is known about the dynamics of cNTS neurons during eating, in part due to the location of the cNTS in the extreme caudal brainstem, which creates technical challenges for performing neural recordings in awake animals. As a result, most of our understanding of cNTS regulation has come from studies of *Fos* expression^11,13,33^, recordings in brain slices^34–37^, or recordings in anesthetized animals^12,38–40^. These studies have shown that a large proportion of cNTS neurons respond to GI signals, such as stretch of the stomach or intestines, and that these responses are relayed by the vagus nerve and topographically organized in the cNTS^12,18^.

Nevertheless, the meaning of these observations for behavior remains unclear, in part because anesthetized and ex vivo preparations lack the natural sensory and motor feedback that is present during a meal. Thus, while cNTS neurons can respond to artificial stimuli such as gastric balloon inflation, it remains unknown to what extent these mechanisms are engaged during normal behavior. In this regard, the cNTS receives abundant descending input from the forebrain^41–44^, the function of which is unknown, and it is possible that this descending input reshapes or gates cNTS sensory representations in awake animals, as has been observed in other sensory systems^45–47^.

Addressing these questions requires recording cNTS activity while animals eat. As a first step toward this goal, we and others recently reported population recordings of several cNTS cell types by fiber photometry, which revealed cell-type-specific differences in activity^17,29^.

However, these bulk measurements report the average responses of hundreds of cells and therefore cannot reveal the tuning of individual neurons. Moreover, because highly selective marker genes exist for only a few, rare cNTS cell types^17,48^, the regulation of most cNTS neurons cannot be assessed in any meaningful way using fiber photometry.

Recently, we developed a preparation that enables for the first time single-cell calcium imaging in the cNTS of awake animals^17^. Here, we use this approach to characterize how food ingestion is broadly encoded across glutamatergic neurons in the cNTS. We find that, contrary to current models of meal termination, most cNTS activity during eating is not driven by GI signals that report on cumulative food consumed. Instead, these circuits are synchronized to rapid, pregastric cues that report on the dynamics of behavior.

## Results

### Most cNTS neurons can be progressively activated by gastrointestinal feedback

The majority of cNTS neurons are glutamatergic^48,49^, and this Vglut2+ population includes all of cNTS cell types known to inhibit food intake^17,21–29,38^. We therefore targeted GCaMP6s to these cells by injecting a Cre-dependent AAV into Vglut2-Cre mice and, in the same surgery, implanted an angled GRIN lens above the cNTS for microendoscopic imaging (Fig. 1a-b). In a separate surgery, mice were equipped with an intragastric (IG) catheter so that we could directly infuse nutrients into the stomach. Mice were then habituated to a head-fixed and lower-body restrained imaging preparation^17^ that greatly reduced motion artifacts and enabled stable imaging of the cNTS during ingestion (Fig. 1c).

**Figure 1:**
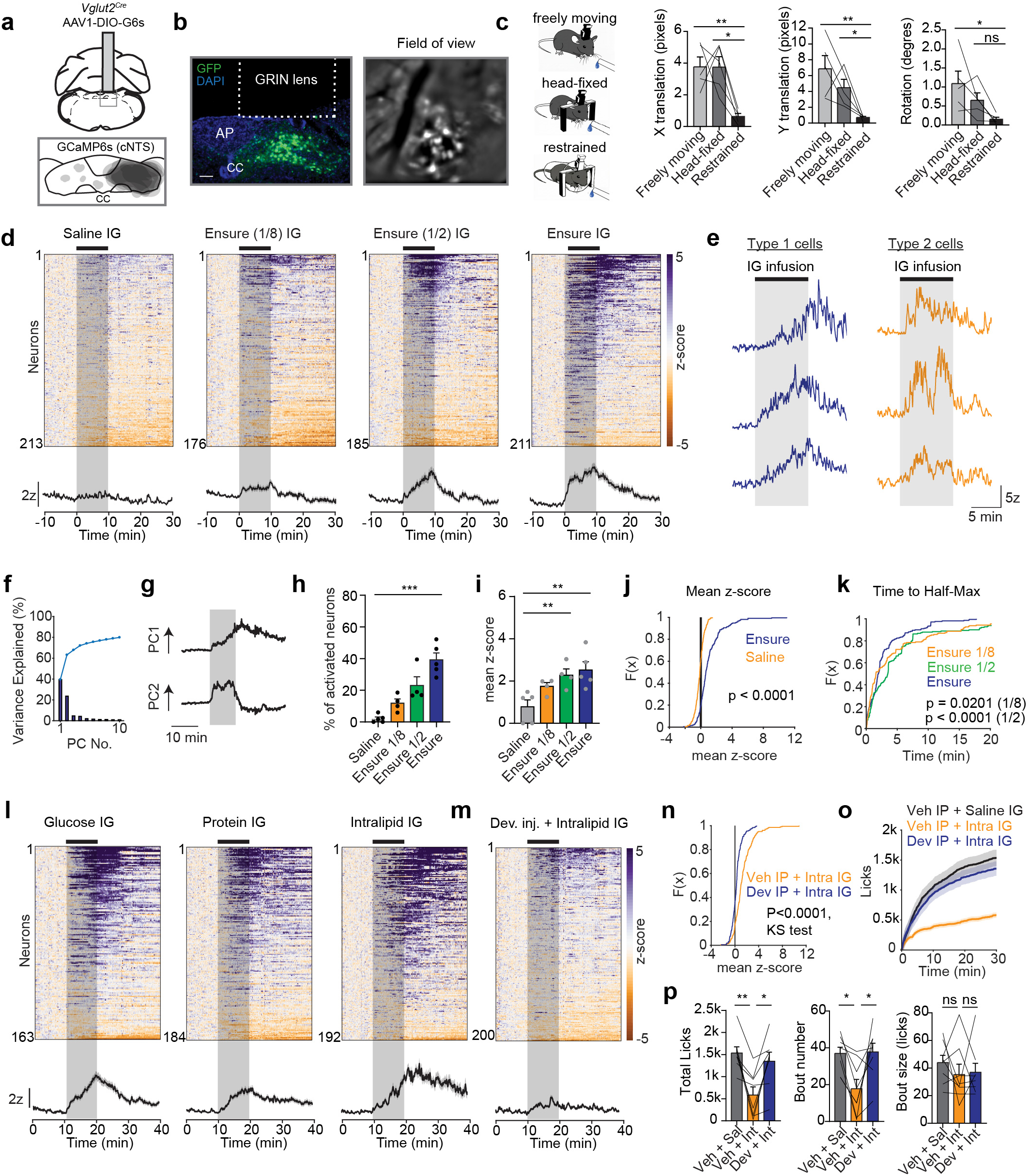
cNTS neuron responses to IG nutrients. a, Top, Schematic of viral targeting strategy showing AAV1-DIO-GCaMP6s injection into the caudal nucleus of the solitary tract (cNTS) of *Vg/ut2c,e* mice. Bottom, Viral expression of GCaMP6s in cNTS neurons below GRIN lens. b, Left, histological verification of GCaMP expression in cNTS neurons and GRIN-lens placement (GFP, DAPI). Right, representative field of view used for microendoscopic imaging. Scale bar= 100 µm. c, Quantification of animal movement across freely moving, head-fixed, and fully restrained conditions (n = 5 mice). Left, X translation (pixels). Middle, Y translation (pixels); Right, rotation (degrees) d, Top, Heatmaps of z-scored calcium activity during lntragastric (IG) infusion of saline or different concentrations of Ensure (1/8 dilution, 1/2 dilution, undiluted; 1 ml). Bottom, Peri-stimulus time histograms (PSTH) across all neurons. e, Individual neuron traces from Type 1 and Type 2 neurons during IG infusion of Ensure. f, Variance explained (%) by the first ten principal components during IG infusion of Ensure. g, PC 1 and 2 for IG infusion of Ensure responses plotted against time. h, Percentage of activated neurons during IG infusion of Ensure at different concentrations. i, Mean z-score (0-30 min) for activated neurons during IG infusion of Ensure at different concentrations. j, Cumulative distribution function (CDF) of mean z-scores (0-30 min) for IG infusion of saline or undiluted Ensure (1ml). k, CDF of time to half-maximal activation for IG infusion of Ensure at different concentrations. I, Heatmaps and PSTHs of isovolumetric and isocaloric IG infusions (1 ml) of glucose (25%), protein (25%, collagen peptides), or lntralipid (10%). m, Heatmaps and mean traces for IG infusion of lntralipid after devazepide injection. n, CDF of mean z-scores (0-30 min) for IG infusion of lntralipid after vehicle or devazepide injection. o, Cumulative licks of a glucose solution (24%) after IG infusion of saline or lntralipid (10%, 0.5 ml) with vehicle or devazepide injection. p, Left, Total licks of a glucose solution after IG infusion of saline or lntralipid with vehicle or devazepide injection. Middle, bout number. Right, bout size (licks). NS, *P<0.05, **P<0.01, ***P<0.001, ****P<0.0001. Data are mean± sem.

Because the cNTS receives extensive direct input from vagal afferents that innervate the gut^34^, we first measured the response to infusion of nutrients into the stomach (Fig. 1d-k and Extended Data Fig. 1a-h). IG infusion of the complete diet Ensure (1 mL, approximating a moderately-sized meal) caused strong activation of a large subset of cNTS neurons (53% activated during the 10 min infusion; Fig.1d-j and Extended Data Fig. 1b). This activation ramped over the course of the ten minute infusion (181 ± 14 seconds, time to 50% of peak activation) and then remained elevated for tens of minutes after the infusion stopped. The response was dose-dependent, with isovolumetric infusions of more concentrated Ensure recruiting more neurons (Fig. 1d,h and Extended Data Fig. 1b) which were more strongly activated (Fig. 1i and Extended Data Fig. 1a) and responded more quickly (Fig. 1k). These responses were not macronutrient specific, as isocaloric and isovolumetric infusions of pure sugar (glucose), fat (Intralipid) or protein (a peptide solution) induced similarly broad activation of cNTS neurons (Fig. 1l and Extended Data Fig. 1e-h). Relative to sugar and protein, the response to isocaloric fat infusion was stronger (3.8 ± 0.6 z for cells activated by Intralipid compared to 2.4 ± 0.2 z and 2.2 ± 0.1 z for glucose and peptides, p = 0.0162) but also more delayed (Extended Data Fig. 1e-g), which is consistent with known differences in the kinetics of digestion and absorption of these macronutrients^50,51^.

Analysis of responsive neurons revealed two broad classes of activated cells (Fig. 1e-g). Type 1 cells ramped during the infusion and then remained elevated for ten or more minutes, whereas Type 2 cells were activated during infusion and then rapidly declined when the infusion stopped (Fig. 1e). Consistently, most of the variance in the population responses could be explained by two principal components which had dynamics that mirrored these two activation patterns (Fig. 1f-g). These responses may relate to the acute and sustained GI signals generated by IG infusion (e.g. gastric accommodation during infusion and sustained nutrient responses afterward). Indeed, we found that many cNTS neurons were activated by either IG infusion of methylcellulose, which induces gastric distension^6,52^, or injection of the hormones CCK and GLP-1, which are released from the intestine in response to nutrients (Extended Data Fig. 1j-p).

We next asked whether blocking the cNTS activation by one of these gut-brain signals would also alter feeding behavior. CCK is particularly important for relaying to the brain information about ingested fat^8,17,40,53,54^, and we found that pretreatment with CCKAR antagonist devazepide greatly reduced cNTS neural activation by IG infusion of Intralipid (Fig. 1m-n and Extended Data Fig. 1q-r). Consistently, pretreatment with devazepide also abolished the ability of an Intralipid preload to reduce subsequent food intake (Fig. 1o-p). Taken together, these data show that IG nutrients activate a large population of cNTS neurons through known mechanisms of gut-brain signaling, and that this activation correlates with changes in feeding behavior.

### cNTS neurons are rapidly activated during oral ingestion

We next investigated how cNTS neurons are regulated during oral food consumption. Mice were fasted overnight and then given access to Ensure for self-paced consumption. Surprisingly, and in contrast to the gradual ramping observed during IG infusion (Fig. 1), we found that most cNTS neurons were activated within seconds during oral consumption (median response latency = 4.7 s vs. 66 s for oral consumption vs. IG infusion, p < 0.0001; Extended Data Fig. 2a-d). Sorting of activated neurons by their response patterns revealed that 70% of these cells were active only during the 10 min ingestion window (suggesting they respond primarily or only to an acute, ingestion-generated signal), whereas 25% were active both during and after ingestion, and 5% were active only after ingestion (suggesting they respond primarily to post-ingestive feedback; Extended Data Fig. 2a). This suggests that, during oral ingestion, most cNTS activity is driven by rapid, likely pregastric, signals associated with eating, rather traditional gut-brain pathways.

To enable direct comparison between oral and IG responses, we performed matched experiments in which mice, on different days, consumed the same food in the same amount and at the same rate by either oral ingestion or IG infusion (Fig. 2a-c). This confirmed that cNTS neurons are activated much more rapidly during oral consumption compared to IG infusion (e.g. median response latency = 4.8 s vs. 97 s for matched oral versus IG consumption of Ensure, p < 0.0001; Fig. 2d-f and Extended Data Fig. 3a-f). In addition, the neural response to oral ingestion decayed much more quickly when consumption stopped (time to 50% decay = 20 ± 2.5 s vs. 87 ± 12 s for oral versus matched IG infusion of Intralipid, p < 0.0001, Extended Data Fig. 3g-j). As result of these differing dynamics, the response to IG infusion was strongly correlated with cumulative food intake, whereas the response to oral ingestion was not (Fig. 2g and Extended Data Fig. 3k-n). Consistently, Intralipid was more satiating when infused into the stomach than when the same amount was consumed by mouth (Fig. 2h-j).

**Figure 2:**
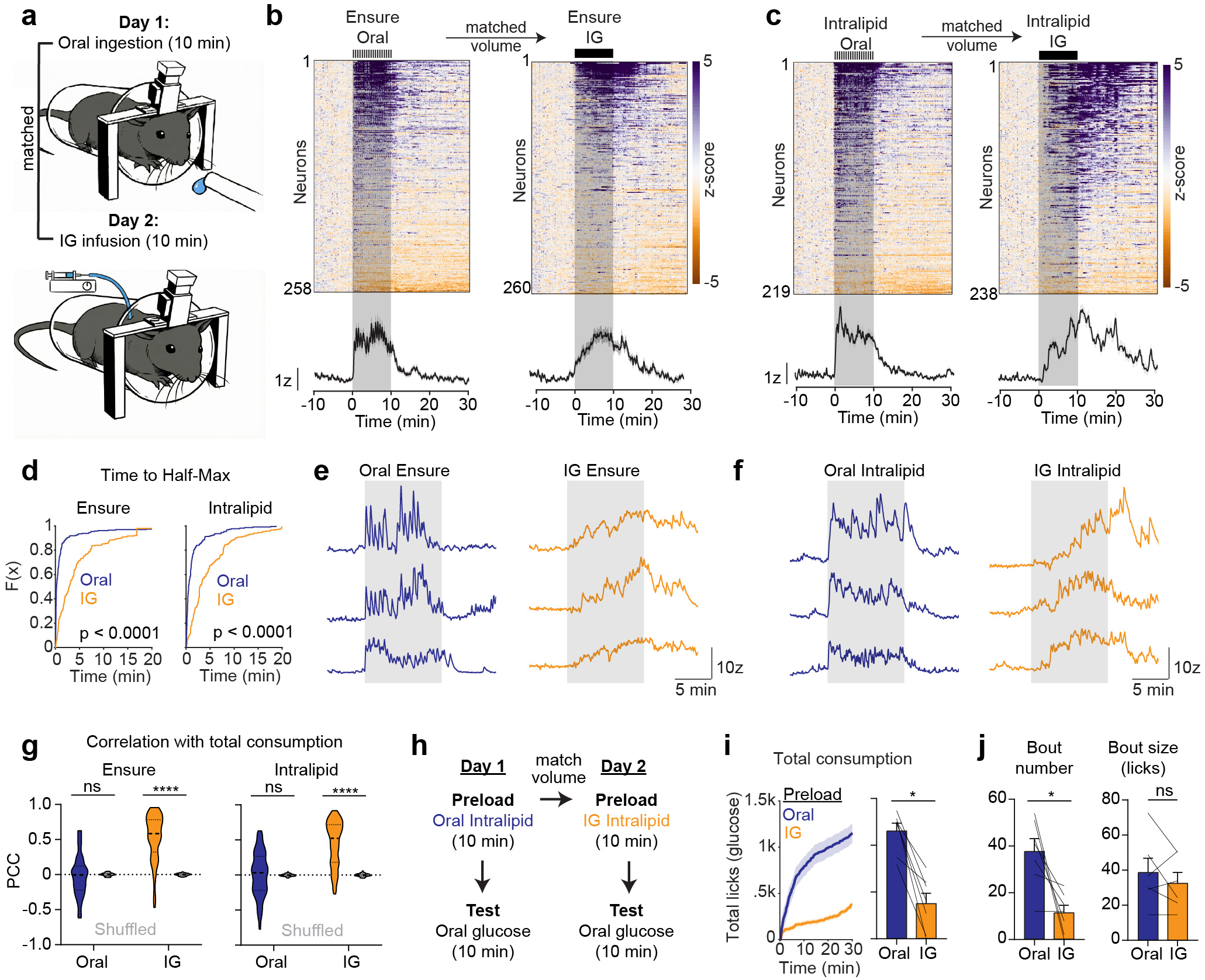
cNTS neurons are rapidly activated during oral ingestion. a, Experimental design for comparing oral ingestion and matched-volume IG infusion of the same nutrients. On Day 1, mice consumed nutrients orally for 10 min. On Day 2, the same matched volume was delivered intragastrically over 10 min. b, Left, Heatmap and PSTH for all neurons during oral Ensure ingestion (left) and matched-volume IG infusion of Ensure (right). c, Left, Heatmap and PSTH for all neurons during oral lntralipid ingestion (left) and matched-volume IG infusion of lntralipid (right). d, Left, CDF of time to half-maximal activation for oral versus IG Ensure. Right, CDF of time to half-maximal activation for oral versus IG lntralipid. e, Individual neuron traces during oral ingestion of Ensure (left) or matched **IG** infusions of Ensure (right) f, Individual neuron traces during oral ingestion of lntralipid (left) or matched **IG** infusions of lntralipid (right) g, Left, Pearson correlation coefficient (PCC) values comparing cNTS neuron activity with total oral or IG intake for Ensure. Right, PCC values comparing cNTS neuron activity with total oral or IG intake for lntralipid. Real data (color) is compared vs. shuffled controls (gray). h, Experimental design for testing how oral versus IG preloads influence subsequent oral glucose intake. i, Total oral glucose consumption following oral lntralipid ingestion or a matched-volume IG infusion of lntralipid. j, Left, bout number during oral glucose intake after each preload condition. Right, bout size (licks). NS, *P<0.05, **P<0.01, ***P<0.001, ****P<0.0001. Data are mean± sem.

The fact that neural responses were sustained after IG infusion but not after matched oral ingestion is paradoxical, because all food consumed by mouth rapidly reaches the stomach and therefore should elicit the same GI feedback. However, it is possible that oral ingestion alters gut reflexes (e.g. slowing of gastric emptying) in a way that diminishes downstream GI signals. To test this, we allowed mice to consume Intralipid by mouth or gave them a matched IG infusion and then characterized the contents of the GI tract. We found that there was no difference in the amount of Intralipid retained in the stomach (0.42 ± 0.08 g vs. 0.49 ± 0.09 g for oral ingestion vs. IG infusion, p = 0.7766) or in different segments of the small intestine after Intralipid delivery by these two routes (Extended Data Fig. 4a-d). There was also no difference in circulating triglyceride levels at various time points after oral versus IG infusion (51 ± 7 mg/dL vs. 48 ± 4 mg/dL for oral Intralipid vs. matched IG infusion after 30 min, p = 0.8438; Extended Data Fig. 4e). This indicates that the different post-ingestive neural responses we observe are not due to gross differences in food transit. Instead, this suggests that the responsiveness of cNTS neurons to GI feedback is generally reduced during oral ingestion, enabling these neurons to instead track rapid, pregastric signals.

### Oral and GI feedback activate partially overlapping ensembles in the cNTS

The fact that cNTS neurons respond to different signals during oral ingestion and IG infusion raises the question of whether these responses are encoded by the same or different cells (Fig. 3a). We therefore presented mice with sequential oral and visceral stimuli and compared the responses of individual neurons (Fig. 3b).

**Figure 3:**
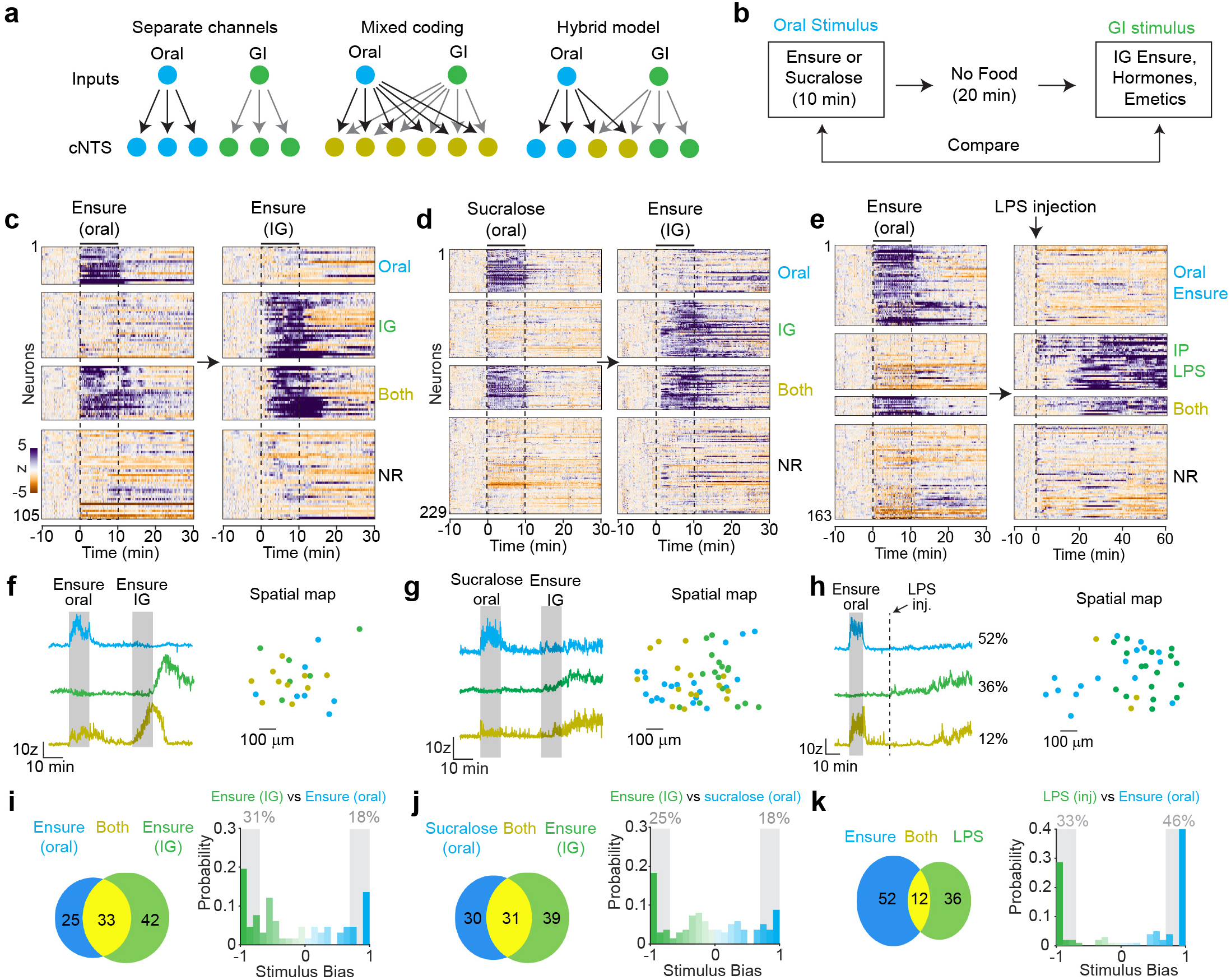
Oral and GI feedback activate partially overlapping ensembles in the cNTS. a, Conceptual models for how oral and gastrointestinal (GI) inputs may be represented in cNTS populations: separate channels, mixed coding, or a hybrid model. b, Experimental design: mice received a 10-min oral stimulus (Ensure or sucralose) followed 20 min later by a GI stimulus (IG Ensure, hormones, or emetics). cNTS neuron responses to oral vs. GI stimuli were compared within the same field of view. c, Heatmaps of responses to oral Ensure ingestion (left) and IG infusion of Ensure (1 ml, right), with neurons classified as responsive to Oral-only, GI-only, Both, or Non-responsive (NR). d, Heatmaps of responses to oral sucralose ingestion (left) and IG infusion of Ensure (right). e, Heatmaps of cNTS neuron activity during oral Ensure ingestion (left) and IP LPS injection (right). f, Left, Individual neuron traces for oral Ensure vs. IG Ensure. Right, spatial map of activated neurons for oral Ensure vs. IG ensure. g, Left, Individual neuron traces for oral sucralose vs. IG Ensure. Right, spatial map of activated neurons for oral sucralose vs. IG ensure. h, d, Left, individual neuron traces for oral Ensure vs. LPS injection. Right, spatial map of activated neurons for oral Ensure vs. LPS injection. i, Left, Venn diagram showing percentage of cells activated by oral Ensure vs. IG Ensure. Right, histogram of stimulus bias index for oral Ensure vs IG Ensure. Stimulus bias= (Stimulus 1 response - Stimulus 2 response)/ (Stimulus 1 response+ Stimulus 2 response). See Methods. j, Left, Venn diagram showing percentage of cells activated by oral sucralose vs. IG Ensure. Right, histogram of stimulus bias indexes for oral sucralose vs. IG Ensure. k, Left, Venn diagram showing percentage of cells activated by oral Ensure vs. LPS injection. Right, histogram of stimulus bias indexes for oral Ensure vs. LPS injection.

We first compared oral versus IG Ensure. Mice were fasted overnight and then allowed to consume Ensure for 10 min, at which point the sipper was removed. Twenty minutes later, mice received an IG infusion of Ensure (1 mL over 10 min) and activity was recorded for an additional twenty minutes. Alignment of single-cell responses (Fig. 3c, f, i) revealed, first, a major subset of cells that respond to oral but not IG Ensure (Type 1 cells, 25% of activated neurons). This shows that many cNTS neurons are specifically tuned to pregastric signals. A second class of cells (Type 2; 42%) were activated by IG infusion of Ensure but not oral ingestion. Thus, these cells can respond to GI feedback, but this is suppressed during oral ingestion. A third group of cells (Type 3; 33%) were activated by both oral and IG Ensure and therefore could theoretically respond to a mixture of pregastric and GI signals during normal ingestion (Extended Data Fig. 5a-d). Consistent with this possibility, we found that while the onset of responses to oral ingestion was equally rapid in Type 1 and 3 cells (Extended Data Fig. 5b), more activation persisted after ingestion ended in Type 3 cells (Extended Data Fig. 5e), reflecting a possible contribution from a delayed GI signal.

Since oral Ensure is not a pure oral stimulus but also contains calories that could influence responses to subsequent IG infusion, we repeated this experiment by comparing IG Ensure with oral sucralose, which is sweet but has no post-ingestive effect (Extended Data Fig. 5f-h). We found that oral ingestion of sucralose and Ensure activated a similar percentage of cNTS neurons (58% vs. 61% for Ensure vs. sucralose), indicating that calories and GI feedback are not required for the acute cNTS response to ingestion (Fig. 3d, g, j). When we sorted these sucralose-activated neurons by their responses to IG Ensure, this revealed again three populations of responsive cells (Fig. 3d) in similar proportions to the previous experiment (Fig. 3c). However, in this case, there was no difference between Type 1 and 3 cells in the activity that persisted after sucralose consumption ended (Extended Data Fig. 5i), which may be due to the fact that sucralose does not have calories. Taken together, these data show that most cNTS neurons respond rapidly to pregastric cues, but that, in a subset of these cells, the duration of these responses may be modulated by post-ingestive feedback.

Much research has focused on the distinction between cNTS neurons involved in satiety and those involved in nausea or sickness^17,22,25,38,55,56^, but these ideas have never been tested by neural recordings in awake animals. We therefore asked to what extent aversive and nutritional stimuli modulate the same versus different cNTS cells. Mice were allowed to consume Ensure (10 min) before being challenged with an injection of either LiCl (an emetic) or LPS (which induces a sickness-like state^56–58^) (Fig. 3e and Extended Data Fig. 6a-g). We found that there were again neurons specifically tuned to either oral or visceral (in this case, aversive) stimuli, as well as some cells that responded to both (Fig. 3h, k and Extended Data Fig. 6b-c). However, compared to the preceding experiments that examined two ingestion-related stimuli, there was noticeably less overlap between the cells that responded to oral Ensure versus aversive agents. We quantified this difference by calculating a stimulus bias index that reflects a neuron’s relative response for one stimulus versus another on a scale from −1 to +1. This revealed more strongly tuned cells when we compared ingestive and aversive stimuli than when we compared ingestive stimuli to each other (e.g. mean absolute value of stimulus bias index = 0.85 ± 0.03 for oral Ensure-LPS injection versus 0.65 ± 0.04 for oral-IG Ensure, p < 0.0001; Fig. 3c and Extended Data Fig. 6f). We obtained similar results when we calculated the correlation between the neural responses to pairs of oral and visceral stimuli (e.g. p < 0.0001 between oral Ensure-IG Ensure versus oral Ensure-LiCl injection; Extended Data Fig. 6g). Taken together, these data support the notion that many cells in the cNTS are biased to respond to either nutritional or aversive signals, although this demarcation is not absolute.

### cNTS neurons respond to a combination of chemosensory and mechanosensory pregastric cues

We next investigated the nature of the signals responsible for the rapid activation of cNTS neurons during ingestion. To separate ingestive from exterosensory (i.e. cue) responses, mice were trained to consume Ensure in a trial-based format in which each trial was initiated by an auditory cue (1s) followed by a delay (2s) and then brief access (30s) to a small volume of Ensure (∼30 uL, diluted 1:1 in water) (Fig. 4a). Approximately 50% of cNTS cells were rapidly activated during ingestion in this paradigm (Fig. 4b). Among these activated cells, the largest group (72% in trial 2) responded time-locked to the onset of consumption, with activity that ramped over the 5s during ingestion before gradually declining after the Ensure was consumed (Fig. 4b-c and Extended Data Fig. 7a). A second group of cells were activated by the auditory cue that preceded food delivery and then became further activated during consumption (21% of activated neurons), whereas a small population of cells responded only to the cue and then declined to baseline after ingestion commenced (7% of activated neurons; Fig. 4c-e). Thus, most cNTS neurons are activated by oral contact with food, although some cells can also respond to cues that predict food delivery.

**Figure 4.**
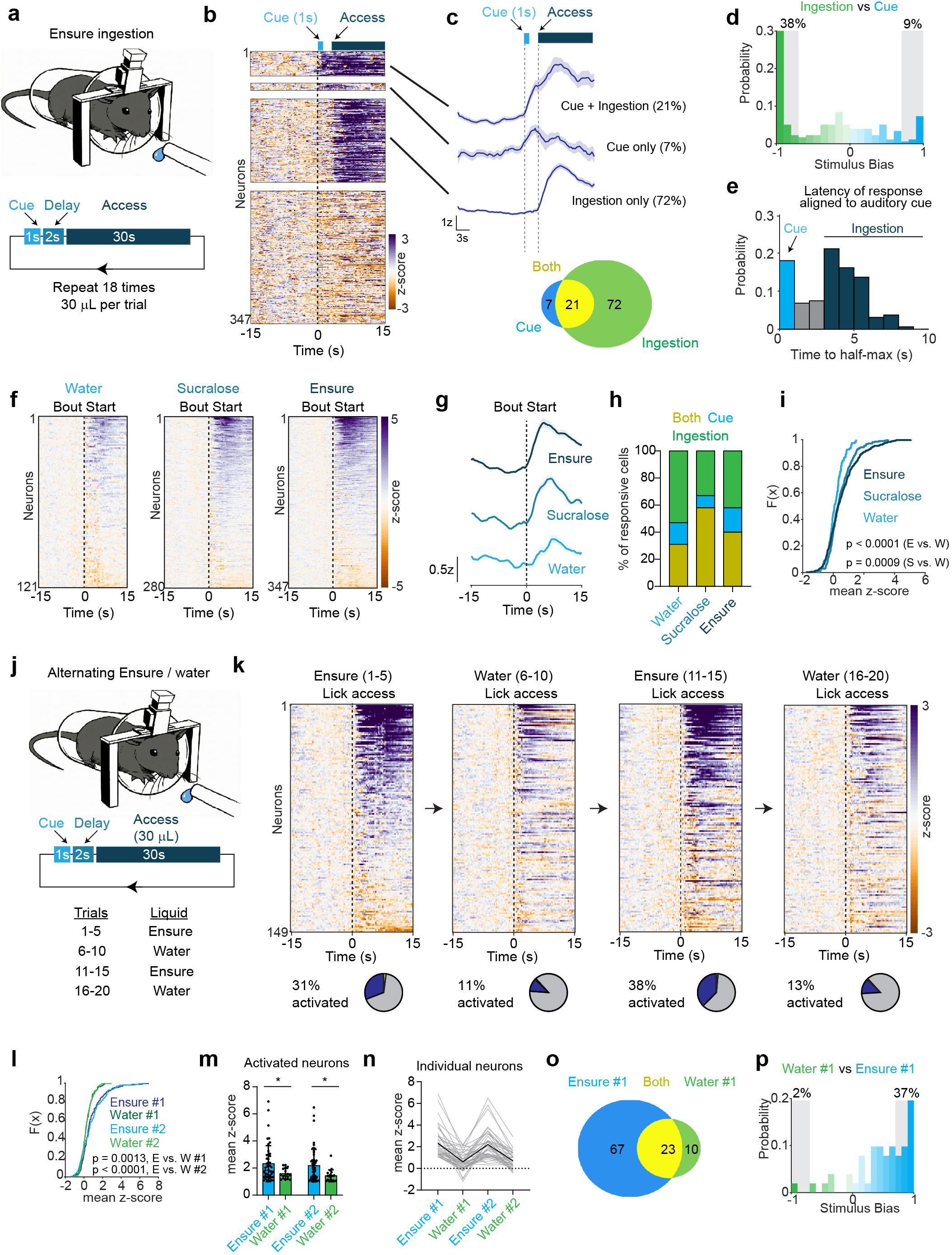
Most cNTS neurons respond rapidly to pregastric cues associated with food ingestion. a, Experimental paradigm for trial-based delivery of Ensure: A 1s auditory cue and a 2s delay preceded 30s access to a liquid drop of Ensure(∼30 µL), which was repeated across 18 trials. b, Heatmap of cNTS neuron activity across one trial, or bout, of Ensure ingestion. Neurons were categorized as responsive to cue and ingestion, cue only, ingestion only, or non-responsive **(NR)**. c, Top, PSTH for each response category aligned to bout start and corresponding percentages. Bottom, Venn diagram showing percentage of cells activated by the cue, ingestion, or both. d, Histogram of stimulus bias indexes for cue (blue) vs. ingestion responses (green). e, Histogram of time to half-maximal activation during one trial of Ensure ingestion. f, Heatmaps of bout-aligned cNTS neuron responses for water, sucralose, and Ensure ingestion. g, PSTH aligned to bout start for water, sucralose, and Ensure ingestion. h, Percentage of neurons classified as responsive to cue only, ingestion only, or both during water, sucralose, and Ensure ingestion. i, CDF of mean z-scores (0-15s) for water, sucralose, and Ensure ingestion across the first 5 trials. j, Experimental paradigm for alternating trials of Ensure and water (four blocks: Ensure 1-5, water 6-10, Ensure 11-15, water 16-20). k, Heatmaps and percentage of activated neurons for each block of Ensure or water ingestion. I, CDF of mean z-scores (0-15s) for cNTS neuron responses to Ensure and water ingestion across alternating blocks. m, Mean z-scores of activated neurons across blocks of Ensure or water ingestion. n, Mean z-score responses of individual neurons tracked across blocks of Ensure or water ingestion. o, Venn diagram showing percentage of cells activated by oral ingestion of Ensure vs. water. p, Histogram of stimulus bias indexes for the first block of Ensure (blue) and water ingestion (green). NS, *P<0.05, **P<0.01, ***P<0.001, ****P<0.0001. Data are mean± sem.

These ingestion-triggered responses did not depend on a specific macronutrient, as we observed similar activation during consumption of pure glucose, Intralipid, or sucralose (Extended Data Fig. 7e), whereas responses to water ingestion were weaker (Fig. 4f-i) despite similar licking behavior (Extended Data Fig. 7f). To understand how individual cells respond to food versus water, we performed experiments in which the access to Ensure or water was interleaved across trials within the same session (Fig. 4j). This confirmed that Ensure consumption recruited more cells than water (38% vs. 13%, second round of presentations) and caused greater mean activation per cell (2.2 ± 0.2 z vs. 1.4 ± 0.1 z for Ensure versus water, second round of presentations, p = 0.0207, Fig. 4k-n), despite no significant differences in licking behavior (Extended Data Fig. 7g). Of note, most of the cells that responded to water ingestion also responded to Ensure (Fig. 4o-p), indicating that few cells are tuned to water specifically. Analogous experiments comparing ingestion of glucose and Intralipid revealed that the population responses to sugar and fat consumption were more similar to each than the responses to Ensure and water (PCC = 0.34 ± 0.02 for glucose-Intralipid compared to 0.16 ± 0.03 for Ensure-water, p < 0.0001; Extended Data Fig. 7h-o), suggesting that most cells respond to food ingestion rather than the specific food identity.

In all the experiments above, the mouse was allowed to freely initiate consumption. To rule out the possibility that these differential neural responses to food and water are secondary to differences in behavior, we performed neural recordings in response to intraoral infusions, which allows us to precisely control the rate, timing, and duration of oral fluid ingestion (Fig. 5a). Mice were given four brief (5 second) infusions of either water, sucralose, or Ensure, each separated by one minute. Consistent with results from free licking, the percentage of responsive neurons progressively increased as the fluid was changed from water to sucralose to Ensure (e.g. 37% to 62% to 76% of cells responding in the first bout). There was also an increase in the magnitude of responses among the activated neurons, with stronger average responses to Ensure and sucralose than water (e.g. 2.9 ± 0.4 z vs. 5.4 ± 0.5 z for water vs. sucralose 15-30s during the first bout, p < 0.0001; Fig. 5b-f and Extended Data Fig. 8a-c). Interestingly, these stronger magnitude responses were specific to the period immediately after fluid ingestion (e.g. 15-30s after the start of each trial, Fig. 5e), which resulted in a ramping activation throughout the session following consumption of sucralose and Ensure but not water (Fig. 5b). Taken together, these data indicate that, relative to water, food tastes and/or nutrients increase cNTS activation by (1) recruiting more cells throughout the session, and (2) increasing the duration of activation of those cells, so that it persists immediately after ingestion stops.

**Figure 5:**
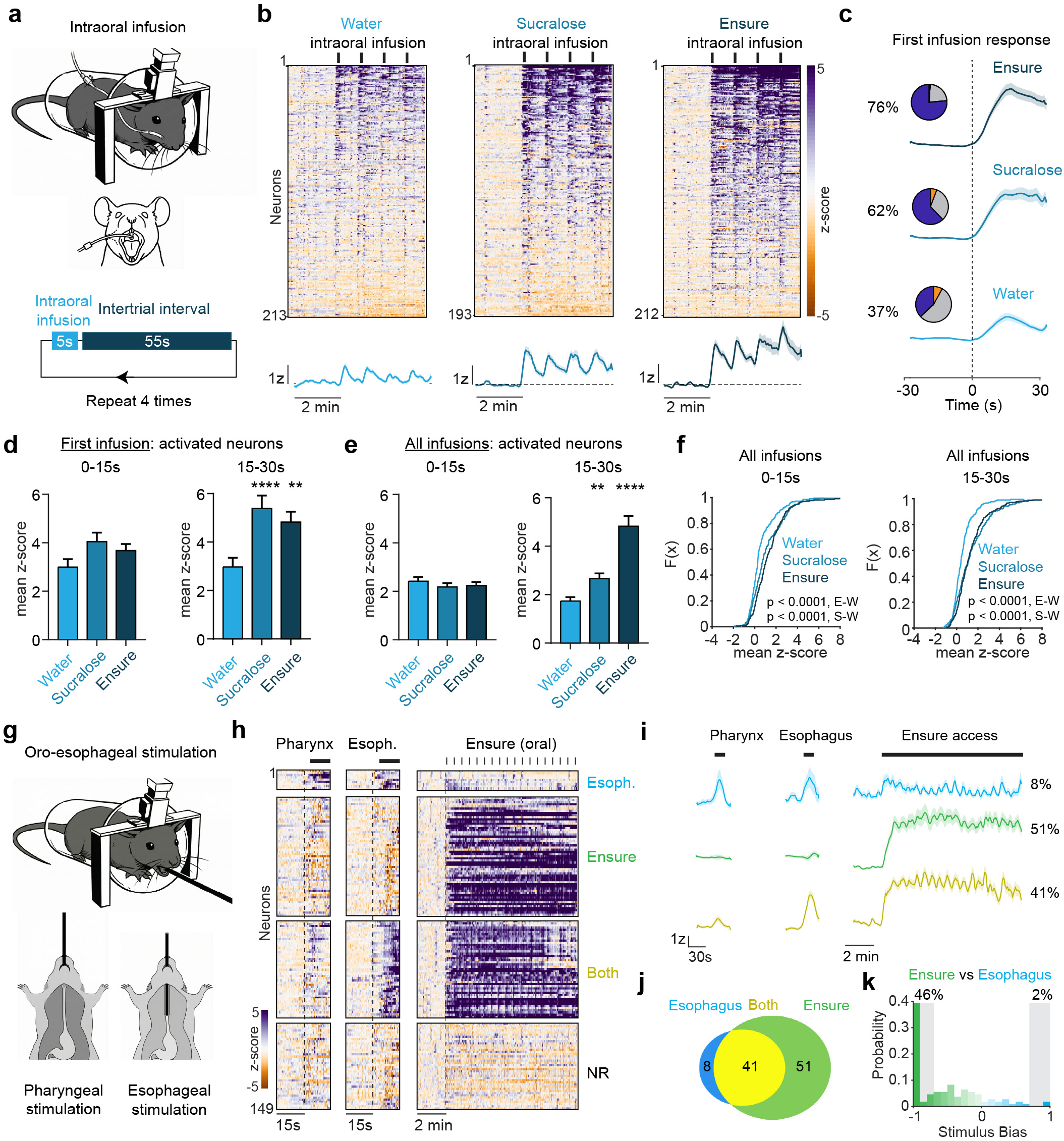
cNTS neurons respond to mechanosensory and chemosensory signals in the oral cavity. a, Schematic of intraoral infusions during cNTS imaging. Mice received four 5s intraoral infusions separated by 55s intertrial intervals. b, Heatmaps of cNTS neuron activity during intraoral infusions of water (left), sucralose (middle), or Ensure (right). lntraoral infusions are indicated by black tick marks. Bottom, PSTH aligned to start of intraoral infusions for water, sucralose, and Ensure. c, Left, pie charts showing the proportion of activated (purple), inhibited (orange), and non-responsive (gray) neurons during the first intraoral infusion for Ensure, sucralose, and water. Right, PSTH of activated neurons aligned to infusion onset for each tastant. d, Mean z-score (0-15s and 15-30s) for activated neurons during the first infusion of water, sucralose, or Ensure. e, Mean z-score (0-15s and 15-30s) for activated neurons across all infusions of water, sucralose, or Ensure. f, CDF of mean z-scores for activated neurons across all infusions of water, sucralose, and Ensure for the 0-1 Ss (left) and 15-30s (right) windows. g, Schematic of pharyngeal (left) and esophageal (right) stimulation during cNTS imaging. h, Heatmaps of cNTS neuron activity aligned to pharyngeal distension (left), esophageal distension (middle), and oral Ensure ingestion (right). Neurons are categorized as esophagus-responsive (blue), Ensure-responsive (green), both (orange), or non-responsive (NR). i, PSTH aligned to stimulus onset of neurons responsive to pharyngeal, esophageal, or oral Ensure stimulation. j, Venn diagram showing percentage of cells responsive to esophageal stimulation (blue), Ensure ingestion (green), or both (yellow). k, Histogram of stimulus bias indexes for esophageal stimulation (green) vs. Ensure ingestion (blue). NS, *P<0.05, **P<0.01, ***P<0.001, ****P<0.0001. Data are mean± sem.

In addition to these chemosensory signals, the volume of fluid ingested can be detected by monitoring distension of the pharynx and esophagus, which is sensed by cranial afferents that innervate the throat and project to the caudal brainstem^20^. To investigate this, we used an oral gavage tube to mechanically stimulate the pharynx or esophagus and then compared these responses to those of Ensure ingestion in the same session (Fig. 5g). We found that many cNTS neurons were activated by distension of the pharynx (20%) or esophagus (36%) (Fig. 5h and Extended Data Fig. 8d). This activation was coincident with tube insertion and then rapidly declined when the tube was removed (Fig. 5i). Subsequent consumption of Ensure recruited a broad population of activated cNTS neurons (67%), which included almost all cells that responded to isolated esophageal distension (83%) (Fig. 5j-k and Extended Data Fig. 8e-g). This indicates that esophageal distension is likely one component of the cNTS response to fluid ingestion. However, the fact that most Ensure responsive cells were not responsive to esophageal stretch (Fig. 5j-k) indicates that other signals are involved.

### GI feedback is largely dispensable for rapid cNTS responses to ingestion

Given that most of the cNTS response to ingestion is too fast to be driven by gut feedback, we designed two experiments to assess whether GI signals are required for these rapid dynamics. First, we tested whether CCK, which is required for cNTS activation in response to IG fat (Fig. 1 and Extended Data Fig. 1) is also required for the cNTS activation in response to orally ingested fat. Remarkably, we found that it was not, as pretreatment with the CCKAR antagonist devazepide had no effect on any aspect of the response to oral fat ingestion, including the percentage of activated cells (30 ± 10% vs. 27 ± 9 % after vehicle vs. devazepide injection, p = 0.9510), the strength of their activation (2.1 ± 0.2 z vs. 2.0 ± 0.3 z after vehicle vs. devazepide, p = 0.9470), their latency to respond or the duration of their response after Intralipid consumption (Extended Data Fig. 8h-n). Consistent with this, devazepide pretreatment also had no effect on the amount of Intralipid consumed by mouth (Extended Data Fig. 8o-q). This was in contrast to IG Intralipid, where devazepide pretreatment caused a dramatic increase in subsequent food consumption (Fig. 1o-p). Thus, the cNTS uses different signals to detect ingested fat when it is consumed by mouth versus when it is infused into the stomach, and CCK becomes dispensable during oral ingestion.

Many gut-brain signals of food ingestion are transmitted by the vagus nerve^1,2,5,6,11,59^. We therefore asked whether gut-innervating vagal afferents are necessary for cNTS responses to food ingestion. To test this, we took advantage of the fact that our imaging preparation is unilateral (i.e. we image only one hemisphere of the cNTS per mouse) and the two nodose ganglia project preferentially to their ipsilateral cNTS^60–62^. For this reason, a unilateral, cervical vagotomy on the same side as the imaging implant will greatly reduce vagal input to the imaged cells, while at the same time leaving most of the contralateral vagal pathway intact. This allows us to block vagal input without causing the disruptions to GI function that accompany bilateral subdiaphragmatic vagotomy and confound the interpretation of behavioral experiments^63–65^.

We first measured cNTS responses, in vagus-intact animals, to Ensure consumption and CCK injection. We used CCK as a control because gut-brain signaling by CCK is known to require the vagus nerve^59,66,67^. We then subjected animals to a vagotomy in which we transected the ipsilateral vagus just below the level of the superior laryngeal nerve (Fig. 6a-b and Extended Data Fig. 9a), which eliminates all vagal input from the viscera (including the stomach, intestines, and lower esophagus) but has minimal effect on vagal innervation of the oropharynx and upper esophagus^68–70^. After animals recovered from surgery, we then recorded again cNTS responses to consumption of Ensure or a CCK injection.

**Figure 6:**
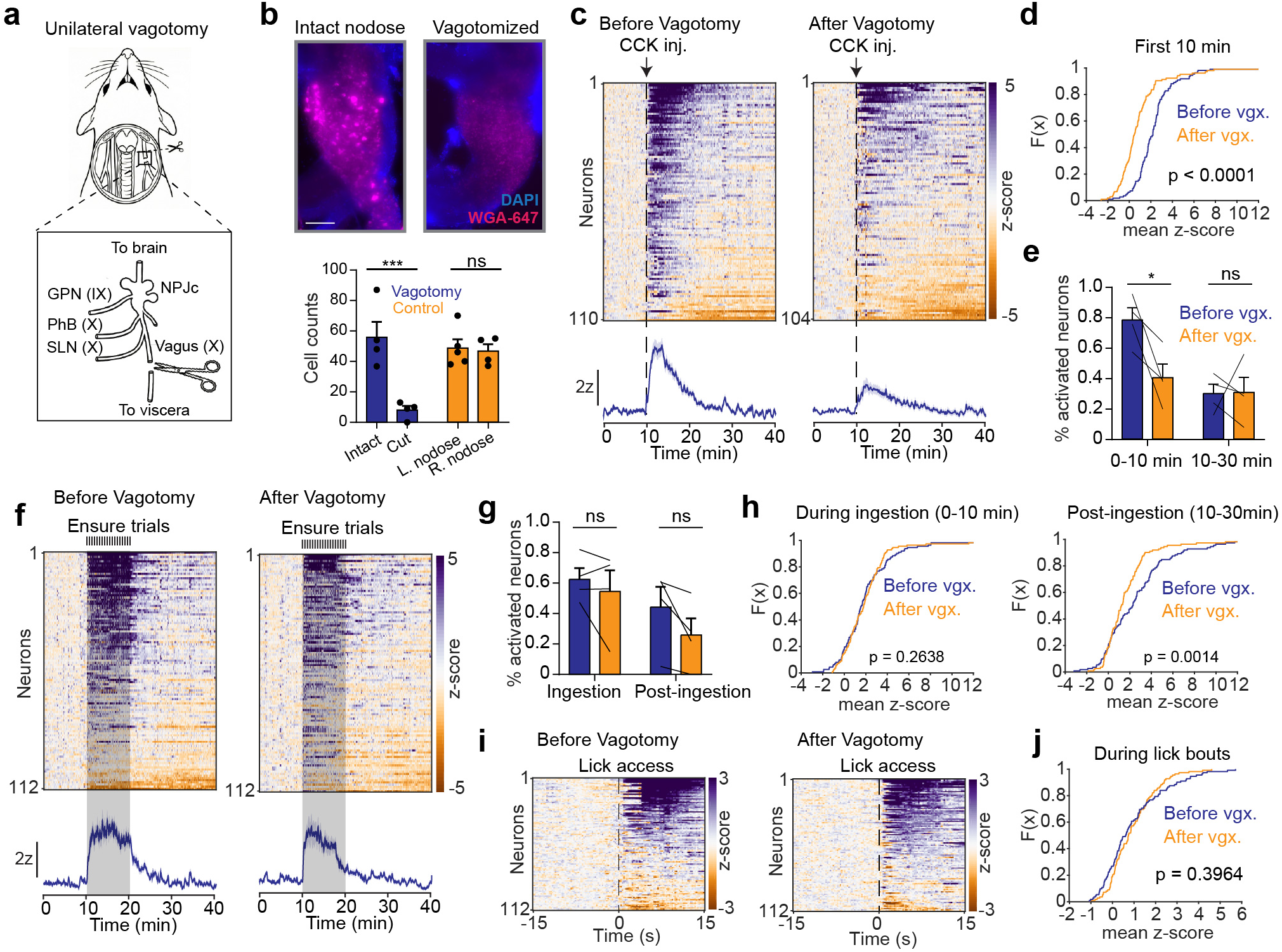
Rapid cNTS activation during ingestion does not require gut-innervating vagal afferents. a, Schematic of unilateral cervical vagotomy targeting the central branch of the vagus nerve (X) innervating the viscera while preserving the glossopharyngeal nerve (GPN; IX), pharyngeal branch of the vagus (PhB; X), and the superior laryngeal nerve (SLN; X). These nerves are connected to the nodose-petrosal-jugular complex (NPJc), which innervate the cNTS in the brain. b, Top, histology for retrograde WGA-647 labeling of nodose ganglion neurons before and after unilateral vagotomy. Bottom, quantification of cells in the nodose ganglion labelled with WGA-647 for mice after vagotomy, or control animals. Scale bar= 500 µm. c, Heatmaps of cNTS neuron activity during subcutaneous CCK injection before and after vagotomy. Bottom, PSTH aligned to injection for all neurons. d, CDF of mean z-scores (0-10 min) for CCK injections before and after unilateral vagotomy. e, Percentage of activated neurons after CCK injection during early (0-10 min) and late (10-30 min) periods, before and after vagotomy. f, Heatmaps of cNTS neuron activity during Ensure ingestion trials before and after vagotomy. Bottom, PSTH aligned to Ensure access for all neurons. g, Percentage of neurons activated during ingestion (0-10 min) and after ingestion (10-30 min) h, CDF of mean z-scores during ingestion (0-10 min) and post-ingestion (10-30 min). i, Heatmaps of cNTS neuron activity during the first five trials of Ensure ingestion before and after vagotomy. j, CDF of mean z-scores (0-1Ss) during lick bouts before and after vagotomy. NS, *P<0.05, **P<0.01, ***P<0.001, ****P<0.0001. Data are mean± sem.

As expected, unilateral vagotomy greatly reduced the cNTS responses to CCK injection, including the percentage of activated cells and the overall strength of activation (78 ± 8 % vs. 40 ± 9 % activated cells, before vs. after vagotomy, p = 0.0151; Fig. 6c-e and Extended Data Fig. 9b). The fact that the blockade was not complete is presumably due to the fact that some vagal afferents project to the contralateral cNTS^60–62^. However, and in striking contrast to the CCK results, vagotomy had minimal effect on most responses to oral ingestion of Ensure (Fig. 6f-j and Extended Data Fig. 9c-e). For example, there was no difference in the percentage of activated cells (62 ± 7 % vs. 55 ± 14 %, before vs. after vagotomy, p = 0.8693) or the strength of their activation (p = 0.6254) over the course of the 10 min Ensure access (Fig. 6g and Extended Data Fig. 9e). There was also no difference in the strength of the seconds-timescale cNTS responses that occurred at the start of each lick bout (Fig. 6i-j and Extended Data Fig. 9f). We did detect a small reduction in vagotomized animals in the persistence of cNTS neuron activation after Ensure access was removed (Fig. 6h), which is consistent with other data (Fig. 3c and Fig. 5b) suggesting that GI input may modulate the duration of pregastric responses. In addition, we found that cNTS neurons were activated even more rapidly by oral ingestion following vagotomy (Extended Data Fig. 9g-j), suggesting that these circuits may become more tuned to pregastric signals after the partial loss of GI feedback. Taken together, these data show that two canonical GI signals – CCK and the vagus nerve – are dispensable for most of the cNTS responses to food ingestion, which instead reflect faster, pregastric signals.

### A PVH to cNTS pathway for rapid responses to ingestion

The fact that most cNTS neuron responses to ingestion do not require gut-innervating vagal afferents raises the question of where these signals come from. The responses to pharyngeal and esophageal distension (Fig. 5g-i) are likely transmitted by branches of the vagal and glossopharyngeal nerves that innervate these structures^20,68^ and which were spared in our vagotomy. However, most cNTS neurons that responded to Ensure ingestion were not activated by esophageal distension, indicating that additional inputs are involved (Fig. 5j). In addition to these cranial nerves, the cNTS also receives extensive descending input from the forebrain^41,44^, raising the possibility that some of these rapid responses may be inherited from other structures. One plausible candidate to provide this input is the paraventricular nucleus of the hypothalamus (PVH), which provides dense glutamatergic input to the cNTS^41,71–73^ and which, like the cNTS, contains neurons whose activity is time-locked to the dynamics of food ingestion^74^.

We first tested whether the PVH neurons that project to the cNTS exhibit ingestion-activated dynamics. A Cre-dependent AAV expressing axon-GCaMP6s was injected into the PVH of Sim1-Cre mice (which labels almost all PVH neurons)^75–77^ and an optical fiber was implanted above the cNTS to record axonal calcium by photometry (Fig. 7a-b). We found that Ensure ingestion triggered rapid activation of PVH terminals that was time-locked to the start of each lick bout and then decayed when the lick bout stopped (Fig. 7c-e). Additionally, PVH terminals were rapidly activated by ingestion of sucralose (0.75 ± 0.18 z, p = 0.0002) but not saline (0.29 ± 0.16 z, p = 0.2424) or when mice performed dry licks at an empty sipper (Fig. 7d-f). Thus, PVH→cNTS axon terminals respond to ingestion with rapid dynamics similar to cNTS neurons, suggesting that PVH input could contribute to these cNTS responses.

**Figure 7:**
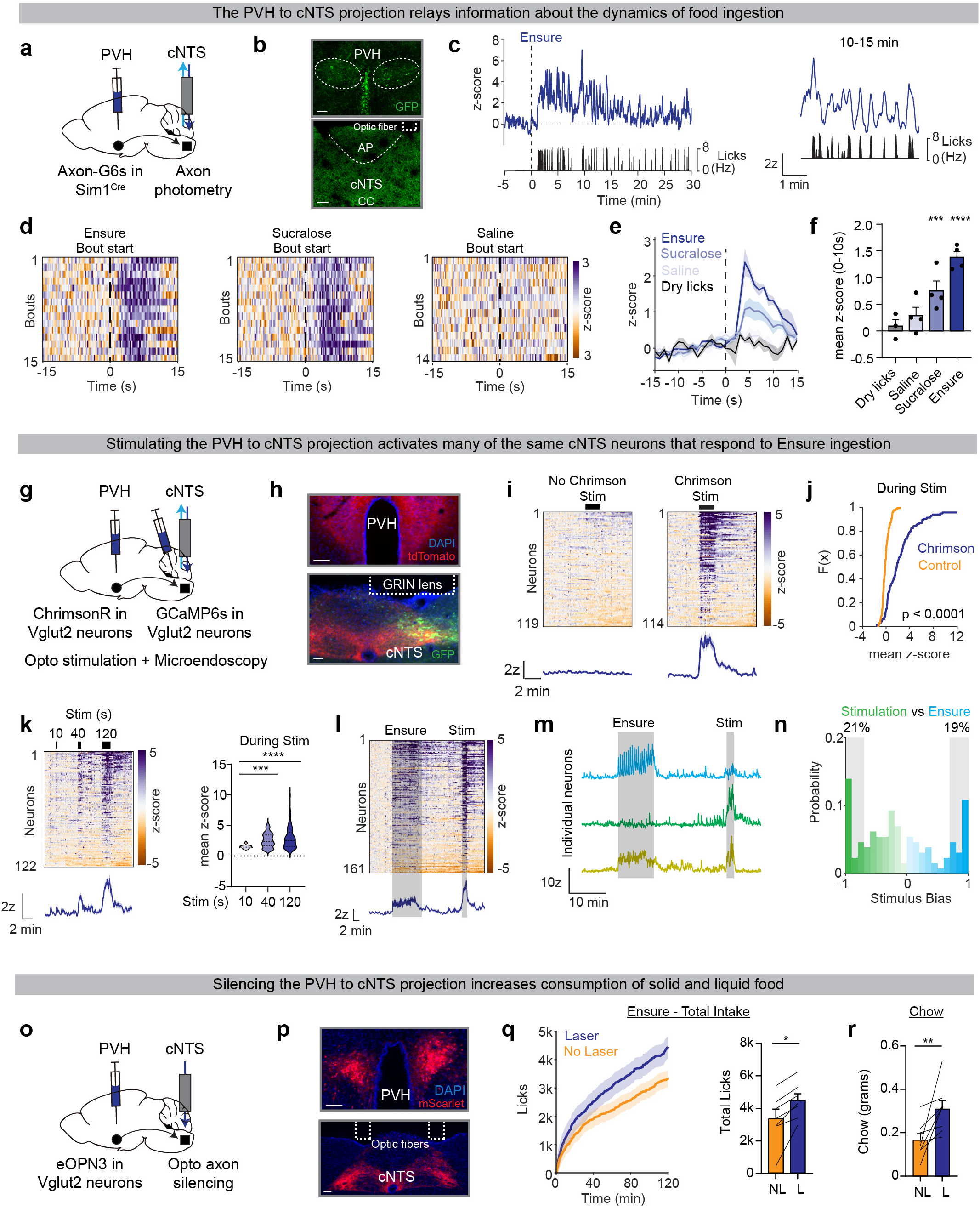
A PVH to cNTS pathway for rapid responses to ingestion. a, Schematic of axon-GCaMP6s expression in PVH Sim1 neurons and axon photometry in cNTS. b, Histological validation of GCaMP labeling in the PVH and cNTS, and optic fiber placement above the cNTS. Scale bar = 100 µm. c, Left, PSTH aligned to Ensure access showing PVH---+cNTS axon activity. Right, zoom in of PSTH from 10 to 15 min. d, Heatmaps of bout-aligned PVH---+cNTS axon activity during Ensure (left), sucralose (middle), and saline ingestion (right). e, PSTH aligned to the first lick for Ensure ingestion, sucralose ingestion, saline ingestion, and dry licking. f, Mean z-scores (0-10s) for all lick bouts during Ensure ingestion, sucralose ingestion, saline ingestion, and dry licking. g, Schematic of ChrimsonR expression in PVH Vglut2 neurons and GCaMP6s in cNTS neurons for simultaneous optogenetic stimulation and microendoscopic imaging in the cNTS. h, Histological validation of ChrimsonR expression in the PVH and cNTS, GCaMP expression in the cNTS, and GRIN lens placement in the cNTS. Scale bar= 100 µm. i, Heatmaps of cNTS neuron activity during optogenetic PVH terminal stimulation versus no-opsin controls. Bottom, PSTH aligned to stimulation onset. j, CDF of mean z-scores (0-2 min during stimulation). k, Left, heatmap of cNTS neuron responses during 10s, 40s, and 120s PVH stimulation epochs. Right, mean z-score during each stimulation epoch for activated neurons. I, Heatmap of cNTS neuron activity during Ensure ingestion followed by PVH terminal stimulation. m, Individual neuron traces showing neurons preferentially responsive to Ensure ingestion, PVH terminal stimulation, or both. n, Histogram of stimulus bias indexes for Ensure ingestion (blue) and PVH terminal stimulation (green). o, Schematic of optogenetic silencing: eOPN3 expression in PVH Vglut2 neurons for silencing of PVH---+cNTS axons. p, Histological verification of eOPN3-mScarlet in the PVH and cNTS, and fiber placement above the cNTS. Scale bar= 100 µm. q, Left, cumulative licks of Ensure during open-loop optogenetic silencing of PVH terminals or no laser trials. Right, quantification of total licks. r, Chow intake during open-loop optogenetic silencing of PVH terminals or no laser trials. NS, *P<0.05, **P<0.01, ***P<0.001, ****P<0.0001. Data are mean± sem.

We next measured how PVH input modulates cNTS neurons in vivo. To do this, we designed an experiment in which we could stimulate PVH terminals while simultaneously imaging calcium dynamics in cNTS cell bodies. A Cre-dependent AAV expressing the red-shifted opsin ChrimsonR was targeted to the PVH of Vglut2-Cre mice and, in the same surgery, GCaMP6s was targeted to the cell bodies of neurons in the cNTS (Fig. 7g-h). We then installed a GRIN lens above the cNTS so that we could image local cell bodies while stimulating PVH axons using an nVoke microscope^78^.

We first showed that stimulation of PVH axons in the cNTS caused rapid activation of a large subset of cNTS cells in mice expressing ChrimsonR but not in controls (Fig. 7i-j and Extended Data Fig. 10a-b). This activation scaled with the duration of axon stimulation, as longer stimulation (from 10 to 40 to 120 s) resulted in the progressive recruitment of more cNTS cells (e.g. 20% to 48% to 61% of cells activated) and stronger activation during stimulation or after laser offset (Fig. 7k and Extended Data Fig. 10c-f). To determine whether the same cNTS cells respond to PVH input and natural ingestion, we allowed mice to lick Ensure for 10 min, then removed access for 20 min and briefly stimulated PVH axons. There was extensive overlap between the cells that responded to Ensure and PVH stimulation (47% of cells responded to both), and their responses were significantly correlated (p < 0.0001, Fig.7l-n and Extended Data Fig. 10g-l). This suggests that PVH input contributes to the natural activation of cNTS during ingestion, although other inputs are likely also involved.

To test the functional importance of this PVH input for behavior, we targeted the inhibitory opsin eOPN3 to the PVH of Vglut2-Cre mice and, in the same surgery, installed a bilateral optical fiber above the cNTS for silencing (Fig. 7o-p). Of note, terminal silencing with eOPN3 blocks synaptic transmission from the targeted axons without impairing firing of distant cell bodies^79^, and therefore allows for functional characterization of individual pathways even when the axon projections are highly collateralized. We found that silencing of PVH to cNTS axons increased consumption of both Ensure and chow, but not water (chow consumed = 0.17 ± 0.03 g vs 0.31 ± 0.04 g, without and with laser, p = 0.0078; Fig. 7q-r and Extended Data Fig. 11a, f-g, j-k), indicating that, like many cNTS cell types^17^, this pathway specifically suppresses food consumption. In contrast, no effect of laser stimulation was observed in control animals that expressed mCherry (Extended Data Fig. 11b-e, h-i, l-m). Taken together, these data show that (1) the PVH relays a lick-triggered signal of ingestion to the cNTS, (2) this pathway activates many of the same neurons that respond to natural ingestion, and (3) this pathway is necessary for restraining consumption of food but not water.

## Discussion

The question of why we stop eating is one of the fundamental problems in physiology. Traditionally, this process has been explained in terms of negative feedback from the stomach and intestines, which gradually intensifies as a meal progresses, tracks cumulative food consumed, and is integrated in meal termination circuits within the cNTS. Here, we have tested this idea by recording for the first time the activity of individual neurons across the cNTS while animals eat. Contrary to current models, we found that most cNTS neurons do not track GI feedback during eating and instead are rapidly activated before any significant amount of food reaches the gut. This rapid feedback is time-locked to ingestion, arises from the mouth and throat, and reflects a combination of mechanical, gustatory and nutritive pregastric signals. These pregastric responses do not depend on canonical gut-brain pathways, including the vagus nerve, although GI feedback likely modulates their duration. We show that one source of this rapid, pregastric input is descending projections from the PVH, which track ingestion dynamics and are required for satiation. These findings challenge the prevailing view that meal termination is driven primarily by gut-brain signals that report on cumulative food consumed. Instead, our data suggest that the caudal brainstem makes extensive use of pregastric signals that track the dynamics of ingestion in order to anticipate when eating should stop.

### cNTS responses are dominated by pregastric signals during ingestion

An unexpected finding from this study is that most cNTS responses to food consumption appear to be driven by pregastric rather than GI cues. This conclusion is based on nine lines of evidence. (1) Most ingestion-activated cNTS neurons respond coincident with, or within a few seconds after, the start of eating (Fig. 4b-e). This timescale precedes any significant food accumulation in the gut^80^. (2) Many cNTS neurons respond to a conditioned auditory cue that precedes the start of eating and predicts food delivery (Fig. 4b-e). (3) The neural response to IG infusion is, on average, much slower than the response to volume, rate and calorie-matched oral consumption of the same food (Fig. 2d and Extended Data Fig. 3a-b). This confirms that GI responses are, indeed, slow and therefore cannot explain the fast dynamics we observe during normal eating. (4) In addition to having a faster onset, the responses to oral consumption have a faster offset than the responses to matched IG infusion (Extended Data Fig. 3g-j), which again suggests different mechanisms. (5) The response to IG infusion is highly correlated with the total amount of food infused, whereas the response to oral consumption is uncorrelated with total intake (Fig. 2g and Extended Data Fig. 3k-l). This result does not make sense if the same signals are driving cNTS activity. (6) Oral ingestion activates many cNTS neurons that do not show any response to IG infusion of nutrients (Fig. 3c-d). Thus, these cells are tuned to pregastric cues and their activation during normal ingestion cannot be attributed to GI signals. (7) Many cNTS neurons are activated by consumption of the non-caloric sweetener sucralose, which has little or no post-ingestive effect (Fig. 3d). (8) Responses to oral consumption are mostly preserved after loss of key gastrointestinal feedback, including notably severing of gut-innervating vagal afferents (Fig. 6f-j) or blocking the hormone CCK (Extended Data Figure 8h-n). (9) Oral sucralose ingestion causes activation of cNTS neurons that persists for several minutes after the end of ingestion (Extended Data Fig. 5i), whereas IG sucralose has no effect. This is important because it reveals that an oral, gustatory stimulus alone can cause responses that are sustained into the post-ingestive period without any involvement from gastrointestinal feedback.

While these observations imply a major role for pregastric signals in driving cNTS dynamics during eating, we did find evidence for an effect of GI feedback in modulating the duration of these responses. For example, we found that whether a neuron responds to IG Ensure predicts the duration of its response to oral Ensure (Extended Data Fig. 5e), even though the rapid onset of those oral responses is not due to GI feedback. We likewise found that, during intraoral infusions, the presence of nutrients increases the duration of activation that persists immediately after the infusion stops (Fig. 5d-f). Similarly, we found that removal of gut-innervating vagal afferents selectively decreases the cNTS activation that persists after the end of oral Ensure consumption (Fig. 6h). All these findings suggest that GI signals exert their force, in part, by modulating the persistence of neural responses that are initiated by pregastric cues. This modulatory effect occurs on a timescale of seconds to a few minutes. In addition to these modulatory effects, we also identified a small percentage of cells (5% during Ensure free licking, Extended Data Fig. 2a) that have a slow onset activation and may be driven entirely by bona fide post-ingestive signals.

### Rapid cNTS activation involves both chemosensory and mechanical signals from the oropharynx and esophagus

Our data suggest that both mechanical and chemical signals from the mouth and throat are involved in the rapid activation of cNTS neurons during eating. Regarding the former, we found that direct distension of the oropharynx or esophagus activates many of the same cNTS cells that respond to eating (Fig. 5j), suggesting that distension is one component of the rapid response to ingestion. Consistently, we found that ingestion of water, which lacks calories and has minimal taste, is sufficient to activate many cNTS neurons (Fig. 4f-g). Nevertheless, we observed much stronger and broader activation when animals ingested nutritive or sweet solutions compared to water (Fig. 4f-p), and these differences persisted even when we precisely matched the ingestion dynamics using an intraoral catheter (Fig. 5a-f). We also found that food consumption caused broader activation than isolated esophageal stimulation (Fig. 5j). These data suggest that there is a baseline response in the cNTS to mechanical signals from the mouth and throat during ingestion, which is then amplified by chemosensory cues associated with food.

How these chemosensory cues become associated with food is an important subject for future investigation. We found that ingestion of a sweet tastant, even when non-nutritive, was sufficient to acutely activate more cNTS neurons than ingestion of water (Fig. 4f-i). This is presumably because sweet taste is associated with nutrient-dense foods, but whether this association in the cNTS is learned or innate is unknown. On the other hand, ingestive responses were stronger and more sustained when an actual nutrient-dense solution (Ensure) was consumed (Fig. 5b,e). This ability of cNTS neurons to rapidly distinguish between nutritive and non-nutritive sweet solutions could involve several mechanisms, including (1) direct nutrient sensing in the mouth^81–83^, (2) modulation of pregastric responses, in real time, by nutrient sensing in the intestine, and (3) a slower learning process in which post-ingestive nutrients feed back to reinforce an association between orosensory cues and caloric value^84–86^. Regarding the last possibility, classic experiments using sham feeding showed that an important component of satiation is pregastric and remains intact even when all GI feedback is lost^87–89^. This residual satiety was hypothesized to be driven by orosensory cues that had become associated, through learning, with a food’s nutrient content^87^. Our discovery of widespread cNTS responses to orosensory cues associated with food is consistent with this behavioral data.

### Gut feedback is reduced, and becomes partly dispensable, during oral ingestion

Signals from the stomach and intestine are thought to be indispensable for satiety, yet we found that cNTS responses to this GI feedback are reduced when food is consumed by mouth. For example, we showed that many cNTS responses to nutrients, particularly fat, are more sustained when they are delivered directly to the stomach compared to when they are ingested orally, even when the food is delivered in the same amount. Moreover, the activation of cNTS neurons by fat depends on CCK only when the fat enters the body through an IG catheter (Fig. 1m and Extended Data Fig. 1q-r). Similarly, we identified many individual neurons that are strongly activated by IG infusion but not at all by matched oral ingestion, which is paradoxical since all food consumed by mouth reaches the stomach. While there are clearly some cNTS neurons that are tuned only to GI signals, our data suggest that most cNTS cells respond weakly to traditional GI feedback during normal ingestion, at least under our imaging conditions.

There are several possible explanations for these observations. One possibility is that oral ingestion causes changes in gut reflexes that minimize the production of the GI signals associated with food ingestion. We were unable to detect differences in gastric emptying or lipid absorption between oral and IG infusion (Extended Data Fig. 4), but it is possible that other reflexes are differentially engaged depending on the ingestion route. We previously identified one rare cell type that responds only to GI feedback (GCG neurons) and, in this case, there was no difference in the neural response to oral versus IG infusion of the same food^17^, which argues against this explanation. However, it is possible that other cell types have different sensitivity to these signals.

A second possibility, which is not mutually exclusive with the first, is that oral ingestion modulates the intrinsic sensitivity of cNTS neurons to signals from the gut. This could involve GABAergic interneurons in the cNTS, which are anatomically positioned to gate vagal input to neighboring glutamatergic cells through feedforward inhibition^90,91^ and which have been shown to shape the responsiveness of glutamatergic cNTS cells to vagal input in anesthetized animals^12^. This would have parallels to other motivational systems, such as the dopamine system, in which GABAergic interneurons function to blunt the response to expected rewards^92^. The effect of such context-dependent gating would be to shift the neural responses to ingestion forward in time, so that pregastric cues, which precede and anticipate the bodily effects of ingestion, can be used to rapidly guide behavior. This would be adaptive in the context of meal termination, because digestion and absorption are slow (e.g. gastric emptying requires hours^93,94^), and satiety must therefore be generated in anticipation of the physiologic effects of eating. Indeed, such anticipatory regulation has now been observed for a number of important cell types that control appetite^17,95–98^. It will be important in the future to investigate how exactly these anticipatory signals are generated and used to guide behavior.

## Methods

All experimental protocols were approved by the Institutional Animal Care and Use Committee of the University of California, San Francisco, following the National Institutes of Health guidelines for the Care and Use of Laboratory Animals.

### Mouse Strains

Animals were maintained in temperature- and humidity-controlled facilities with 12 h light-dark cycle and *ad libitum* access to water and standard chow (PicoLab 5053. All studies employed a mixture of male and female mice. We obtained wild-type (C57BL/6J, Jackson cat.no.000664), *Vglut2^Cre^*(B6;Slc17a6^tm2(cre)Lowl^/J, Jackson cat.no.016963), and *Sim1^Cre^*(B6; Tg(Sim1-cre)^1Lowl^/J, Jackson cat.no. 006395) mice from Jackson Labs. All transgenic or knock-in mice used in these studies were on the C57BL/6J background.

### Intracranial Surgery

#### General procedures

Animals were anaesthetized with 2% isoflurane and placed in a stereotaxic head frame on a heating pad. Ophthalmic ointment was applied to the eyes and subcutaneous injections of meloxicam (5 mg kg^−1^) and sustained-release buprenorphine (1.5 mg kg^−1^) were given to each mouse before surgery. The scalp was shaved, scrubbed (betadine and alcohol three times), a local anesthetic was applied (bupivacaine 0.25%) and then an incision was made through the midline. A craniotomy was made using a dental drill (0.5 mm). Virus was injected at a rate of 150 nL min^−1^ using a glass pipette connected to a 10 µl Hamilton syringe (WPI), controlled using a Micro4 microsyringe pump controller (WPI). The needle was kept at the injection site for 2 min before withdrawal. Fiber optic cannulas or a GRIN lens were implanted after virus injection during the same surgery, and these were secured to the skull using Metabond (Patterson Dental Supply, 07-5533559, 07-5533500; Henry Schein, 1864477) and Flow-It (Patterson Dental Supply, 07-6472542).

#### Microendoscopy in the cNTS

*Vglut2^Cre^* mice (n = 19) were prepared for imaging by injecting AAV1-CAG-Flex-GCaMP6s-WPRE-SV40 (200 nL; 3 x 10^12^ viral genome copies (vg) mL^-1^; Addgene) into the cNTS (1.3 mm AP; ±0.3 mm ML; −4.3 mm DV relative to the occipital crest with 20° in the AP direction) and installing a GRIN lens (9 x 1 mm length; Inscopix 1050-004596) above the injection site (1.5 mm AP; ±0.37 mm ML; −4.27 mm DV relative to the occipital crest with 20° in the AP direction) in the same surgery. The GRIN lens was slowly lowered into the brain at the rate of 3 µm/second with the Siskiyou MC1000e-1 controller. After at least two weeks of recovery from the lens implantation surgery, mice were anesthetized, and head bars were affixed to the skull with Metabond. A baseplate (Inscopix 100-000279) was placed above the lens and affixed with Metabond. When mice were not being used for imaging experiments, a baseplate cover (Inscopix 100-000241) was attached to prevent damage to the GRIN lens. In subsequent surgeries, mice were equipped with an intragastric or intraoral catheter.

#### Fiber photometry recordings in the cNTS

*Sim1^Cre^* (n = 4) mice were prepared for photometry recordings by injecting AAV1-DIO-hSyn-DIO-axon-GCaMP6s (200 nL; 1.2 x 10^13^ viral genome copies (vg) mL^-1^; Addgene) into the PVH (−0.7 mm AP; ±0.25 mm ML; −4.7 mm DV). In the same surgery, an optical fiber (Doric Lenses, MFC_400/430- 0.48_6.5mm_MF2.5_FLT) and sleeve (Doric Lenses, SLEEVE_BR_2.5) were installed above the ipsilateral cNTS (1.3 mm AP; ±0.3 mm ML; −4.15 mm DV relative to the occipital crest with 20° in the AP direction). Mice were allowed to recover for a minimum of three weeks before the first photometry experiment.

#### Integrated optogenetics and calcium imaging in cNTS

*Vglut2^Cre^* (n = 4) mice were prepared for dual optogenetic and imaging experiments by injecting AAV5-FLEX-Syn-ChrimsonR-tdTomato (100 nL; 8.5 x 10^12^ viral genome copies (vg) mL^-1^; Addgene) bilaterally into the PVH (−0.7 mm AP; ±0.25 mm ML; −4.7 mm DV) and AAV1-CAG-Flex-GCaMP6s-WPRE-SV40 (200 nL; 3 x 10^12^ viral genome copies (vg) mL^-1^; Addgene) into the cNTS (1.3 mm AP; ±0.3 mm ML; −4.3 mm DV relative to the occipital crest with 20° in the AP direction). In the same surgery, a GRIN lens (9 x 1 mm length; Inscopix 1050-004596) was installed above the cNTS (1.5 mm AP; ±0.37 mm ML; −4.27 mm DV relative to the occipital crest with 20° in the AP direction). Mice were allowed to recover for a minimum of three weeks before the first imaging experiment.

#### Optogenetics in cNTS

*Vglut2^Cre^* (n = 14) mice were prepared for optogenetic experiments by injecting AAV1-SIO-eOPN3-mScarlet (100 nL; 8 x 10^12^ viral genome copies (vg) mL^-1^; Addgene) or AAV1-hSyn-DIO-mCherry (100 nL; 8 x 10^12^ viral genome copies (vg) mL^-1^; Addgene) bilaterally into the PVH (−0.7 mm AP; ±0.25 mm ML; −4.7 mm DV). In the same surgery, a dual fiber optic cannula (Doric, DFC_200/245-0.37_6.5mm_DF0.8_FLT) and sleeve (Doric Lenses, SLEEVE_BR_2.5) were installed above the cNTS (1.3 mm AP; 0mm ML; −3.95 mm DV relative to the occipital crest with 20° in the AP direction). Mice were allowed to recover for a minimum of three weeks before optogenetic experiments.

### Intragastric catheter surgery

Mice were anesthetized deeply with 2% isoflurane and surgical sites were shaved and cleaned with betadine and ethanol. Subcutaneous injections of meloxicam (5 mg/kg) and sustained-release buprenorphine (1.5 mg/kg) were given to each mouse prior to surgery. A midline abdominal skin incision was made, extending from the xyphoid process about 1.5 cm caudally, and a secondary incision of 1 cm was made between the scapulae for externalization of the catheter. The skin was separated from the subcutaneous tissue using blunt dissection, such that a subcutaneous tunnel was formed between the two incisions along the left flank to facilitate catheter placement. A small incision was made in the abdominal wall, and the catheter (Intech, C30PU-RGA1439) was pulled through the intrascapular skin incision and into the abdominal cavity using a pair of curved hemostats. The stomach was externalized using atraumatic forceps and a purse string stitch was made in the middle of the forestomach using 7-0 non-absorbable Ethilon suture. A puncture was then made in the center of the purse string, and the end of the catheter was inserted and secured by the purse string suture. For the gastric implant, 2-5 mm of catheter end was fixed within the stomach.

At the end of the surgery, the abdominal cavity was irrigated with 1mL of sterile saline and the stomach was replaced. The abdominal incision was closed in two layers, and the catheter was sutured to the muscle layer at the interscapular site. The interscapular incision was then closed and the external portion of the catheter capped using a 22-gauge PinPort (Instech, PNP3F22). Mice received Baytril (5 mg/kg) and warm saline at the end of surgery and were allowed to recover for one week prior to imaging experiments.

### Intraoral catheter surgery

Mice were anesthetized deeply with 2% isoflurane and surgical sites were shaved and cleaned with betadine and ethanol. Subcutaneous injections of meloxicam (5 mg/kg) and sustained-release buprenorphine (1.5 mg/kg) were given to each mouse prior to surgery. An incision of 2 cm was made near the right scapulae for externalization of the catheter. The skin was separated from the subcutaneous tissue using blunt dissection, such that a subcutaneous tunnel was formed between the incision and the right side of the face to facilitate catheter placement. One end of the catheter (Instech, C30PU-RGA1439) was attached to a 23 gauge needle (BD, 305145), which was used to puncture the buccal mucosa between the incisors and molars. The catheter attached to the needle was then pulled through the cheek incision and the subcutaneous tunnel on the right scapula using curved hemostats. The catheter was pulled through until the PE spool collar (sutured to the catheter) was snug against the buccal mucosa, preventing the catheter from escaping the oral cavity. The needle was removed from the catheter and excess catheter tubing was trimmed. The scapular incision was then closed and the external portion of the catheter capped using a 22-gauge PinPort (Instech, PNP3F22).

### Unilateral vagotomy

Under isoflurane anesthesia, a midline ventral neck incision was made to expose the salivary glands, which were gently separated and retracted laterally. The sternomastoid, sternothyroid, and omohyoid muscles were then retracted to reveal the carotid triangle. The carotid sheath was carefully opened, and the vagus nerve isolated for transection (>2 mm segment removed) below the level of the superior laryngeal branch. The incision was immediately closed with absorbable sutures, and mice were allowed to recover. In rare cases, transient ipsilateral ptosis (Horner’s sign) was observed, likely due to inflammation of the cervical sympathetic trunk; otherwise, the procedure was well tolerated without complications.

All recordings were performed within two weeks after surgery, and post-mortem inspection confirmed no visible nerve regrowth. In a subset of mice, intraperitoneal injection of WGA Alexa Fluor 647 (100 µg per mouse) was performed after experimental completion. One week later (3 weeks after surgery), nodose ganglia were dissected and examined, revealing labeling primarily restricted to the ganglion contralateral to the vagotomy, confirming lasting nerve transection.

### Microendoscopic Imaging

#### Microendoscopic setup

Data was collected using Inscopix nVoke microscopes. Videos were acquired at 8 Hz (20% LED power, 8.0 gain, 2x spatial downsampling) using Inscopix software (Data Acquisition Software v.151; https://support.inscopix.com/support/products/nvista-30-and-nvoke-20/data-acquisitionsoftware-v151). After acquisition, videos were first pre-processed, spatially (binning factor of 2) and temporally (binning factor of 2) downsampled, and motion-corrected using Inscopix software (v.1.7, http://support.inscopix.com/mosaic-workflow). Videos underwent additional motion correction with the Mosaic software (v.1.7, http://support.inscopix.com/mosaic-workflow), which produced estimates of the motion artifacts when mice were freely moving, head-fixed, or head-fixed and restrained (Figure 1). Activity traces for individual neurons were then extracted from these videos using the constrained non-negative matrix factorization (CNMF-E) pipeline in the Inscopix software. After initial CNMF-E segmentation, extracted neurons were manually refined to avoid potential confounding factors from uncorrected motion artifacts, region of interest duplication and over-segmentation of the same spatial components.

#### Behavior

To habituate mice to the imaging setup, we head-fixed mice using a commercial mouse head-fixation system (LabeoTech) before applying additional restraint by placing the animal in a 50 mL conical tube (Fisher Scientific 14-432-22). A disposable fluff underpad (MSC281230) was used to reduce limb movement while ensuring that animals were comfortable under restraint. All mice were initially habituated to the imaging setup by receiving trial-based access to liquid drops of Ensure with the nVoke camera (Inscopix) attached for two sessions (2 hours each) on at least two separate days. At the start of each trial, an auditory cue (Adafruit Large Enclosed Piezo Element, ADA1739) was played before a drop of Ensure (∼30 µL) was delivered using a water dispenser with lickometer (LabeoTech).

On the day of the experiment, mice were head-fixed and restrained using the method described above and given 10 min for habituation, with the Inscopix camera turned on. Mice were given stimuli only after an additional 10 min baseline recording period.

For hormone injection experiments (Extended Data Figure 1), mice received subcutaneous injections near the scapular region with compounds at concentrations based on previously published reports: CCK octapeptide 10 or 30 μg/kg (Bachem), exendin-4 100 μg/kg (Tocris), devazepide 1 mg/kg (R&D Systems), LiCl 84 mg/kg (Sigma), and LPS 500 µg/kg (Sigma). All of these compounds were dissolved in saline (0.9%) while devazepide was dissolved in 5% DMSO, 5% Tween-80 in saline (vehicle solution for devazepide). All compounds were injected at the volume of 10 µL/gram of mouse body weight.

For intragastric infusion experiments (Figure 1), solutions - saline (0.9%), glucose (25%), collagen peptides (25%, Sports Research), Intralipid (10%), Ensure (undiluted, 1/2 dilution in water, 1/8 dilution in water), methylcellulose (1%, Bio-Techne), sucralose (0.8g/L, dissolved in saline) - were delivered using a syringe pump (Harvard Apparatus 70-2001) over 10 min. Before mice were placed on the head-fix stage for habituation, the intragastric catheter was attached to the syringe pump using plastic tubing and adapters (AAD04119, Tygon; LS20, Instech). For experiments involving sequential administration of methylcellulose and CCK, methylcellulose (1%) was first infused over 10 min before a 20 min waiting period, and then CCK (30 μg/kg) was injected subcutaneously near the scapular region. For IG infusions of Intralipid with CCKAR blockade, mice were given a subcutaneous injection of vehicle or devazepide (1 mg/kg). After 5 min, the mice were given an IG infusion of saline or Intralipid 10% (0.5 mL) over 10 min and microendoscopic imaging was performed for an additional 20 min.

For free-access Ensure experiments (Extended Data Figure 2), mice were food-deprived prior the experiment. Mice were given a lickometer containing undiluted Ensure for self-paced consumption over 10 min before the lickometer was taken away. Microendoscopic imaging was performed for an additional 20 min after lickometer removal.

For volume-matched Ensure or Intralipid experiments (Figure 2), mice were food-deprived prior to the experiment. On day 1, mice were given access to a lickometer containing Ensure (1/2 dilution in water) or Intralipid (10 %) for 10 min before removal and an additional 20 min of microendoscopic imaging. Liquid drops of Ensure or Intralipid (∼30 µL) was dispensed every 33 s for a maximum of 18 trials per animals. Two days later (day 2 of experiment), overnight fasted mice were given an IG infusion of Ensure or Intralipid over 10 min at the same volume that each animal individually consumed on day 1. In summary, mice received the same volume of glucose or Intralipid solution with the same timing on day 1 and 2, through oral ingestion or IG delivery.

For sequential oral and GI stimulus experiments (Figure 3), mice were food-deprived overnight before getting trial-based access to a lickometer containing Ensure (1/2 dilution) or sucralose for 10 min before removal. After a 20 min delay period, mice were given different subcutaneous injections of CCK (30 μg/kg), Exendin-4 (100 μg/kg), LiCl (84 mg/kg), LPS (500 µg/kg), or an IG infusion of undiluted Ensure (1mL over 10 min). For the experiment involving sequential presentation of sucralose and methylcellulose, methylcellulose (1%, 1mL over 10 min) was infused 40 min after lickometer removal.

For trial-based lickometer experiments (Figure 4), mice were food-deprived overnight or ad libitum fed (Ensure only). Mice were given a lickometer containing Ensure (1/2 dilution in water), Intralipid (10 %), glucose (12%), sucralose (0.8g/L, dissolved in saline), or water for 10 min before lickometer removal and additional 20 min of microendoscopic imaging. After a 1s auditory cue and a delay period of 2s, a liquid drop (∼30 µL) was dispensed and there was a 30s delay until the start of the next trial. Animals received a maximum of 18 trials.

For alternating access to Ensure and water (Figure 4), mice were food-deprived overnight before getting trial-based access to lickometers containing Ensure or water. Animals were allowed to consume Ensure for trials 1-5 and 11-15, and water for trials 6-10 and 16-20. Liquid drops of Ensure or water (∼30 µL) were dispensed every 33s. For alternating access to glucose and Intralipid (Extended Data Figure 7), animals were allowed to consume glucose (24%) for trials 1-5 and 11-15, and Intralipid (20%) for trials 6-10 and 16-20.

For intraoral infusion experiments (Figure 5), mice were food-deprived overnight before receiving 4 infusions of Ensure, sucralose (0.8g/L, dissolved in saline), or water. Each infusion (∼30 µL over 5s) was delivered one minute apart using a syringe pump (Harvard Apparatus 70-2001). Before mice were placed on the head-fix stage for habituation, the intraoral catheter was attached to the syringe pump using plastic tubing and adapters (AAD04119, Tygon; LS20, Instech).

For pharyngeal and esophageal stimulation experiments (Figure 5), mice were food-deprived overnight before a 24-gauge oral gavage tube (FST, 18061-24) was inserted and held in the pharynx (∼1.5 cm from mouth) for 5s before removal. Five minutes later, the same tube was inserted and held in the upper esophagus (∼2.5 cm from mouth) for 5s before removal.

Finally, after 5 minutes, mice were given a lickometer containing Ensure for trial-based consumption. Liquid drops of Ensure were dispensed every 33s for 10 min.

For oral Intralipid ingestion experiments with CCKAR blockade (Extended Data Figure 8), mice were food-deprived overnight and then given a subcutaneous injection of vehicle or devazepide (1 mg/kg). After 5 min, mice were given a lickometer containing Intralipid (10%) for trial-based consumption. Liquid drops of Intralipid were dispensed every 33s for 10 min. Finally, microendoscopic imaging was performed for an additional 20 minutes after lickometer removal.

For imaging experiments before and after unilateral vagotomy (Figure 6), microendoscopic imaging was first performed during CCK injections or oral Ensure consumption prior to vagotomy. On day 1, mice received subcutaneous injections near the scapular region with CCK (10 μg/kg). On day 2, mice were food-deprived overnight and then given a lickometer containing Ensure for trial-based consumption as described previously. Next, mice underwent unilateral vagotomy surgery where the vagus nerve (ipsilateral side of the imaging implant) was transected near the cervical region. After one week of recovery, microendopic imaging was again performed during CCK injections or oral Ensure consumption as described above.

For PVH terminal stimulation experiments (Figure 7), mice were food-deprived overnight before receiving 2 min of optogenetic stimulation via the nVoke microscope (5 mW; 20 Hz with 10 ms pulse width; 2s ON, 3s OFF) and an additional 8 min of microendoscopic imaging. In a separate experiment, mice were given 10s, 40s, and then 2 min of optogenetic stimulation spaced 5 min apart. To compare cNTS responses to PVH terminal stimulation and Ensure ingestion, mice were food-deprived overnight before receiving a lickometer containing Ensure for trial-based consumption. Liquid drops of Ensure were dispensed every 33s for 10 min. After a 20 min waiting period, mice received optogenetic stimulation via the nVoke microscope for 2 min, before an additional 8 min of microendoscopic imaging.

#### Analysis

For each experiment, activity traces for individual neurons were extracted for each mouse and all responses were normalized using the function *z = (C_raw_ - μ)/σ*, where C_raw_ is an output of the Inscopix software, *μ* is the mean *C_raw_* during the 10 min baseline period before stimulus presentation and *σ* is the standard deviation of *C_raw_* during the same baseline period. We defined a neuron as “activated” if the mean z-score was ≥ 1z. Neurons with a mean z-score of < −1z was defined as “inhibited” and a mean z-score between −1z and 1z was defined as “non-responsive”. For most experiments, to calculate the mean z-score during and after ingestion, we computed the mean z-scored changes in activity from 0-10 min for the ingestion period and 10-30 min for the post-ingestion period.

For analysis of motion artifacts (Figure 1), estimates of translation in the X or Y dimension (pixels), or rotation (degrees), were generated by the Mosaic software package during motion correction.

To perform principal component analysis, we used the “pca” function in MATLAB to calculate the variance explained (%) for the first 10 principal components. The PCA scores for the first two PCs were plotted against time (x-axis).

To calculate the time to half-max activation for each cell (Figure 1), we determined the earliest time point during the 30 min trial where the z-scored activity exceeded 50% of the peak z-score across the session.

To analyze responses to IG infusion of methylcellulose and CCK injection (Extended Data Figure 1), we computed the mean z-scored change in activity for each cell during IG methylcellulose infusion (0-10min) and after CCK injection (0-10 min). The 10-minute period prior to each stimulus was used to calculate the mean baseline activity.

To calculate the response latency (Extended Data Figure 2), we used the “sgolayfilt” function in MATLAB to apply the Savitzky-Golay filter (polynomial order = 2, frame length = 5) to activity traces of single cells. We used the “gradient” function in MATLAB on the smoothed data and found the earliest time point where the derivative value was greater than 4 x Standard deviation of the derivative during the 0-10 min baseline period.

To calculate the Pearson correlation coefficient (PCC) for relationships between two variables for each cell (Figure 2 and Extended Data Figure 3), we used the “corrcoef” function in MATLAB with cumulative licks from 0-10 min as an input vector. The z-scored change in activity (compared to the 10 min baseline period) over 0-10 min was used as the other input vector. To calculate the PCC for the shuffled controls, the input vector for food intake was scrambled for each animal.

To calculate the time of peak activation for each cell (Extended Data Figure 2), we computed the mean z-scored change in activity within 30s bins across the entire 30 min trial.

To calculate the time to half-max activation for each cell (Extended Data Figure 3), we determined the earliest time point during the 0-10 ingestion period where the z-scored activity exceeded 50% of the peak z-score.

To calculate the decay constant (Extended Data Figure 3), we determined the earliest time point where the z-scored change in activity decreases below 50% of the value immediately after ingestion/IG infusion ends.

To analyze responses to oral ingestion and various visceral stimuli (Figure 3), we computed the mean z-scored change in activity for each cell during Ensure ingestion (0-10 min), sucralose ingestion (0-10 min), IG infusion of Ensure (0-10 min), IG infusion of methylcellulose (0-10 min), CCK injection (0-10 min), Exendin-4 injection (0-10 min), LiCl injection (0-30 min), LPS injection (30-60 min). The 10-minute period prior to each stimulus was used to calculate the mean baseline activity.

To calculate the stimulus bias index for each cell (Figure 3), we used the mean z-scored change in activity in response to each stimulus to compute: Stimulus bias index = (Stimulus 1 mean z – Stimulus 2 mean z) / (Stimulus 1 mean z + Stimulus 2 mean z). All negative values were set to zero. Therefore, all stimulus bias index values ranged from +1 to −1, where +1 indicates a cell preferentially responding to stimulus 1 and −1 indicates a cell preferentially responding to stimulus 2. To compare the tuning across stimulus pairs (Extended Data Figure 5 and 6), we determined the absolute value for all stimulus bias index values for each pair of stimuli. Cells were defined as “strongly tuned” if their stimulus bias index was less than −0.7 or greater than 0.7.

To calculate the PCC for pairs of stimuli (Extended Data Figure 5 and 6), the z-scored activity traces for both stimuli were used as the input vectors. The “corr” function in MATLAB was used to calculate the PCC for each cell.

For analysis of single-cell responses during lick bouts, the 15 s prior to the start of each lick bout was used to calculate the baseline activity, which was used to calculate the z-score from 0-15s. We defined a neuron as “activated” if the mean z-score was ≥ 1z, “inhibited” if the mean z-score ≤ −1z, or “non-responsive” if the mean z-score was between −1 and 1. We calculated the mean z-scored change in activity for the first 5 trials during Ensure, sucralose, and water ingestion.

To calculate the time to half-max activation for each cell during a lick bout (Figure 4), we determined the earliest time point during 0-15s where the z-scored activity exceeded 50% of the peak z-score across the bout.

To analyze responses to the auditory cue and ingestion (Figure 4), we computed the mean z-scored change in activity after the trial start for the cue (0-3s) and ingestion (3-15s).

To analyze responses to alternating presentations of Ensure and water, or glucose and Intralipid (Figure 4), we calculated the mean z-scored change in activity across trials 1-5 (Ensure #1 or glucose #1), trials 6-10 (water #1 or Intralipid #1), trials 11-15 (Ensure #2 or glucose #2), and trials 16-20 (water #2 or Intralipid #2). The mean z-scored changes were used to calculate the stimulus bias index.

To analyze responses to intraoral infusion (Figure 5), we calculated the mean z-scored change in activity 0-30s for the first infusion or across all four infusions.

To analyze responses to pharyngeal or esophageal stimulation (Figure 5), we calculated the mean z-scored change in activity 0-15s during insertion of the oral gavage tube into the pharynx or esophagus. To compare responses with Ensure ingestion, we calculated the mean z-scored change in activity during Ensure access (0-10 min) or the first bout (0-15s).

To analyze responses to CCK injection or Ensure ingestion before and after vagotomy (Figure 6), we calculated the mean z-scored change in activity 0-10 min and 10-30 min after CCK injection and 0-15s for each Ensure trial. To calculate the time to half-max, we determined the earliest time point during the 0-10 min ingestion period, or 0-15s during the first bout, where the z-scored activity exceeded 50% of the peak z-score.

To analyze responses to PVH terminal stimulation in the cNTS (Figure 7), we calculated the mean z-scored change in activity 0-2 min during optogenetic stimulation and 2-10 min after stimulation offset. To compare the three stimulation durations, we calculated the mean z-scored change in activity 0-10s, 0-40s, and 0-2 min during optogenetic stimulation. To compare PVH terminal stimulation responses with Ensure ingestion, we calculated the mean z-scored change in activity during Ensure access (0-10 min) or the first bout (0-15s). To calculate the PCC between Ensure ingestion and PVH terminal stimulation, the z-scored activity traces for both stimuli were used as the input vectors. The “corr” function in MATLAB was used to calculate the PCC for each cell.

### Fiber photometry

#### Photometry setup

Mice were tethered to a patch cable (Doric Lenses, MFP_400/460/900-0.48_2m_FCM-MF2.5). Continuous 6 mW blue LED (470 nm) and UV LED (405 nm) served as excitation light sources. These LEDs were driven by a multichannel hub (Thorlabs), modulated at 305 Hz and 505 Hz respectively, and delivered to a filtered minicube (Doric Lenses, FMC6_AE(400-410)_E1(450-490)_F1(500-540)_E2(550-580)_ F2(600-680)_S) before connecting through optic fibers (Doric Lenses, MFP_400/460/900-0.48_2m_FCM-MF2.5). GCaMP calcium GFP signals and UV isosbestic signals were collected through the same fibers back to the dichroic ports of the minicube into a femtowatt silicon photoreceiver (Newport, 2151). Digital signals sampled at 1.0173 kHz were then demodulated, lock-in amplified, and collected through a processor (RZ5P, Tucker-Davis Technologies). Data were then collected through the software Synapse (TDT), exported via Browser (TDT), and downsampled to 4 Hz in MATLAB before analysis.

#### Behavior

For all recordings, mice were placed in sound-isolated behavioral chambers (Coulbourn, Habitest Modular System) without water or food access unless otherwise specified. Chambers were cleaned between experiments to remove olfactory cues from previous experiments. Mice were habituated for one night in the chambers before experiments. On a subsequent day, mice were attached to photometry patch cords and habituated to the chambers for a second session. Before each recording, photometry implants on individual mice were cleaned with 70% ethanol using connector cleaning sticks (MCC-S25) and connected to a photometry patch cable immediately afterward. For all photometry experiments, mice were acclimated to the behavior chamber for 20 min with recording before presentation of a stimulus.

For lick response experiments (Figure 7), mice were food-deprived overnight prior to the experiments before receiving access to a lickometer containing the appropriate solution for the entire 30 min session. Solutions were prepared using deionized water at the following concentrations: 0.009 g/mL saline (0.15 M, 0.9%) and 0.8 mg/mL sucralose (in saline). Vanilla Ensure was used directly from the bottle without dilution.

#### Analysis

GCaMP6s calcium responses at 470 nm excitation were normalized to the 405/415 nm “isosbestic” signal using a linear regression model of both signals during the baseline period to generate *F_normalized_* (the fluorescence predicted using the signal obtained with 405/415 nm excitation). Data were analyzed using the function *z = (F_normalized_ - μ)/σ*, where *F_normalized_* is the normalized photometry signal, μ is the mean *F_normalized_* during the baseline period before stimulus presentation and *σ* is the standard deviation of *F_normalized_* during the same baseline period. For example, traces from individual mice, the z-score trace and lick rate were smoothed using a moving average filter with spans of 20 and 5 data points. For data presentation only, plotted mean traces (30 min) were additionally downsampled by a factor of 50 (this was done to decrease the size of each graph).

For analysis of photometry responses time-locked to licking, the 15 s prior to the start of each lick bout was used to calculate the baseline activity. A lick bout was defined as any set of licks that (1) last four or more seconds and (2) are separated from the previous lick bout by at least 20 s. Z-score per bout was calculated as the average z-score in the first 10s of each licking bout from individual animals.

### Behavior Experiments

#### General setup

For all experiments, mice were placed in sound-isolated behavioral chambers (Coulbourn, Habitest Modular System; Med Associates, Davis Rig) without water or food access unless otherwise specified. Chambers were cleaned between experiments to remove olfactory cues from previous experiments. Mice were habituated for one night in the chambers before experiments. On the day of experiments, mice were acclimated to the behavior chamber for at least 10 min.

For experiments involving a fixed IG preload of Intralipid (Figure 1), mice were food-deprived overnight before the experiment was performed in the light phase. After habituation, mice were given an IP injection of vehicle or devazepide (1 mg/kg). After 5 min, the mice were given an IG preload of saline or Intralipid 20% (0.5 mL) over 10 min and then a 10 min waiting period. Finally, mice were given access to a lickometer containing a glucose solution (24%) for self-paced consumption over 30 min.

For matched oral and IG preload experiments (Figure 2), mice were food-deprived overnight before the experiment was performed in the light phase. On day 1, mice were given access to a bottle containing Intralipid 20% for self-paced consumption over 10 min, and the amount consumed was measured. After a 10 min waiting period, mice were given access to a lickometer containing a glucose solution (24%) for self-paced consumption. Two days later, mice were given an IG preload of Intralipid over 10 min, with a volume matched to the amount of Intralipid consumed on day 1. After a 10 min waiting period, mice were given access to the same glucose solution for self-paced consumption.

For experiments involving Intralipid ingestion with CCKAR blockade (Extended Data Figure 8), mice were food-deprived overnight before the experiment was performed in the light phase. After habituation, mice were given an IP injection of vehicle or devazepide (1 mg/kg). After 5 min, the mice were given a lickometer containing Intralipid (20%) for self-paced consumption for 2 hours.

#### Analysis

For licking bout analyses, a licking bout was defined as any set of licks containing at least three licks, in which no inter-lick interval is greater than five seconds. Bout size was calculated as the average number of licks from all lick bouts. The bout number was the total number of bouts across the entire trial.

### Optogenetics

#### Laser parameters

For continuous optogenetic inhibition experiments, the laser was modulated at 20 Hz for a 2 s ON and 3 s OFF cycle with a 10 ms pulse width. Photoinhibition was delivered by a DPSS 473-nm laser (Shanghai Laser and Optics Century BL473-100FC) through a dual fiber optic patch cord (Doric, DFP_200/220/900-0.37_2m_DF0.8-2FC0) at a laser power of ∼5 mW, which was measured at the tip of the patch cable before each day’s experiments.

#### Behavior

All experiments were fully counterbalanced for the order of stimulation and contained both within animal (+/- laser) and fluorophore (mCherry) controls. For all experiments, mice were placed in sound-isolated behavioral chambers (Coulbourn, Habitest Modular System) without water or food access unless otherwise specified. Chambers were cleaned between experiments to remove olfactory cues from previous experiments. Mice were habituated for one night in the chambers before experiments. On a subsequent day, mice were attached to optogenetic patch cords and habituated to the chambers for a second session. On the day of experiments, mice were acclimated to the behavior chamber for 10 min prior to optogenetic manipulation and food/water access.

For experiments measuring Ensure consumption (Figure 7), animals were ad libitum fed and the experiments were performed in the dark phase (after 5 pm). After habituation, animals were given access to a bottle of Ensure for self-paced consumption over 2 hours as they received open-loop silencing or no laser treatment.

For experiments measuring chow consumption (Figure 7), animals were ad libitum fed and the experiments were performed in the dark phase (after 5 pm). After habituation, animals were given access to a pellet of standard chow (PicoLab 5053) for self-paced consumption over 1 hour as they received open-loop silencing or no laser treatment

For experiments measuring Ensure consumption in fasted animals (Extended Data Figure 11), animals were food-deprived overnight, and the experiments were performed in the light phase. After habituation, animals were given access to a bottle of Ensure for self-paced consumption over 1 hour as they received open-loop silencing or no laser treatment.

For experiments measuring water consumption (Extended Data Figure 11), animals were water-deprived overnight, and the experiments were performed in the light phase. After habituation, animals were given access to a bottle of water for self-paced consumption over 1 hour as they received open-loop silencing or no laser treatment.

#### Analysis

To measure chow consumption, the pellet was weighed before and after the experiment. For licking bout analyses in optogenetic experiments, a licking bout was defined as any set of licks containing at least three licks, in which no inter-lick interval is greater than five seconds. Bout size was calculated as the average number of licks from all lick bouts. The bout number was the total number of bouts across the entire trial.

### Histology

Mice were anesthetized under isoflurane and then transcardially perfused with phosphate-buffered saline (PBS, 10 mL) followed by formalin (10%, 15 mL). Brains were dissected, post-fixed in 10% formalin overnight at 4 °C, and switched to 30% sucrose the next day. All tissues were kept in 30% sucrose at 4 °C for overnight cryo-protection and embedded in OCT before sectioning. Sections (50 μm) were prepared with a cryostat and collected in PBS or on Superfrost Plus slides. To visualize fluorescent labelling without staining, sections were mounted with DAPI Fluoromount-G (Southern Biotech) and then imaged using the Nikon Eclipse Ti2-E.

To measure GI contents in Extended Figure 3, mice received an oral or matched-volume IG preload of Intralipid (20%) for 10 min prior to a 20 min delay, before they were anesthetized under isoflurane and then transcardially perfused with phosphate-buffered saline (PBS, 10 mL) followed by formalin (10%, 15 mL). The pyloric sphincter was tied with a polyester string before the stomach was cut from the small intestine. The small intestine was cut into three segments with similar lengths: proximal, middle, and distal intestine. The mass of the stomach or intestinal segments was weighed before and after the contents were emptied, to measure the amount of food inside each region.

To image nodose ganglia after unilateral vagotomies, animals were transcardially perfused before the left and right nodose ganglia were dissected from each mouse. Intact nodose ganglia were collected on Superfrost Plus slides and mounted with DAPI Fluoromount-G (Southern Biotech) before imaging (Nikon Eclipse Ti2-E). Cells labeled with WGA-647 were manually counted using ImageJ.

### Plasma Triglycerides Assay

To measure plasma triglycerides after an oral or IG preload of Intralipid (Extended Data Figure 4), we used a Triglyceride Colorimetric Assay Kit (Cayman Chemical, 10010303). On day 1, mice were food-deprived overnight before receiving a bottle of Intralipid 20% for self-paced consumption over 10 min. Blood was collected 30, 60, and 120 minutes after the start of Intralipid access. On day 2, or two days later, the same animals given a matched-volume IG preload of Intralipid over 10 min before their blood was drawn at the same 30, 60, and 120-min time point after the IG preload start. Plasma samples were isolated for analysis according to the kit’s instructions.

### Statistics

All values are reported as mean ± sem (error bars or shaded areas). Sample size is the number of animal subjects per group, or number of cells when two distributions are compared. In figures, asterisks denote statistical significance: *p < 0.05, **p < 0.01, ***p < 0.001, ****p < 0.0001. Except for two-way analysis of variance (ANOVA), non-parametric tests were uniformly used. P values for paired or unpaired comparisons were calculated using the Wilcoxon Signed Rank test or Mann-Whitney U test. P values for comparisons across multiple groups were calculated using the Kruskal-Wallis test or two-way ANOVA and corrected for multiple comparisons using Dunn’s multiple comparisons test and Šidák’s multiple comparisons test, respectively. To compare two distributions, the Kolmogorov-Smirnov test was used, and the False Discovery Rate (Benjamini-Hochberg procedure) was used to correct for multiple comparisons. No statistical method was used to predetermine sample size. Randomization and blinding were not used. We analyzed fiber photometry data, behavior data, and microendoscopic imaging data with MATLAB (v.R2017b, http://www.Mathworks.com/ products/matlab) scripts found at https://github.com/zackknightlabucsf/zackknightlabucsf.

## Acknowledgements

We thank J. Kuhl for illustrations and members of the Knight laboratory for comments on the manuscript. Some illustrations were made using Nano Banana Pro. This work was supported by National Institutes of Health grants R01-DK106399, R01-DK138127 and R01-DK145100 (Z.A.K.) and F31DK137586 (T.L.). T.L. is a Stanford Science Fellow at Stanford University. X.Y. is a UCSF Discovery Fellow. Z.A.K. is an Investigator of the Howard Hughes Medical Institute.

## Author Contributions

T.L. and Z.A.K. conceived the project and designed experiments. T.L., X.Y., and G.R.L. led and performed experiments. T.L., X.Y., G.R.L., Y.S., H.N., and Z.A.K. analyzed and interpreted data. T.L., X.Y., G.R.L, and J.Y.O. performed microendoscopic imaging experiments. T.L. performed all photometry experiments. T.L., X.Y., G.R.L., and N.S. performed optogenetic and behavioral experiments. T.L performed all photometry and GRIN lens surgeries. L.Q. performed all intragastric surgeries. T.L. performed all intraoral catheter surgeries. J.C.R.G. performed all unilateral vagotomies. X.Y., T.L, and G.R.L. collected and analyzed histology. T.L., X.Y. and Z.A.K. wrote the manuscript with input from all authors.

## Additional information

Correspondence and requests for materials should be addressed to Zachary A. Knight.

## Data availability

The data from this study are available from the corresponding author on reasonable request.

## Code availability

Links to the code used for data analysis are found at https://github.com/zackknightlabucsf/zackknightlabucsf.

---

**Extended Data Figure 1:**
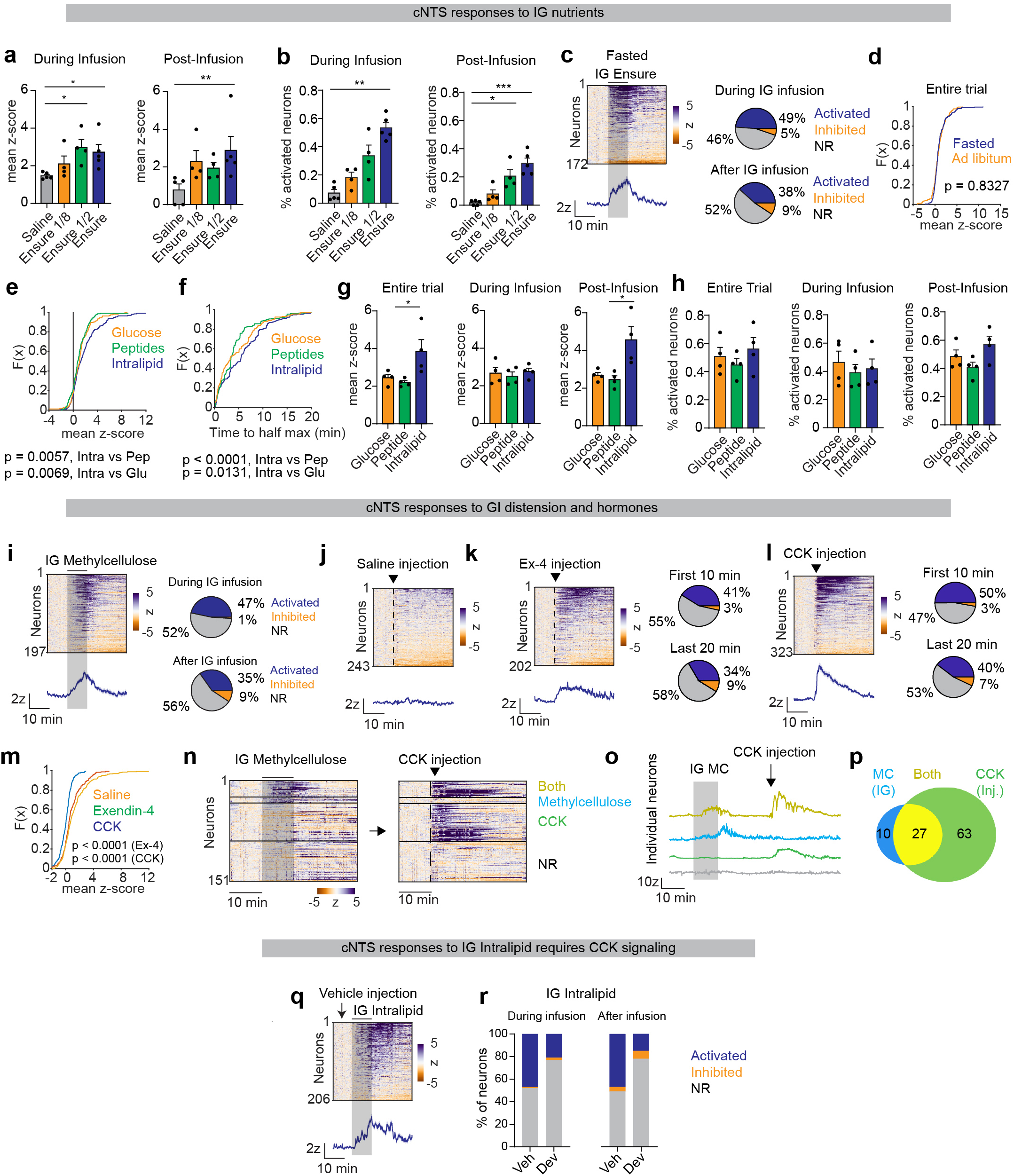
Regulation of cNTS neurons by GI feedback. a, Mean z-scores of activated neurons during IG infusion (0-10 min) and post-infusion periods (10-30 min) for saline, Ensure, glucose, peptides, and lntralipid. b, Percentage of activated neurons during IG infusion (left) and post-infusion (right) of saline, Ensure, glucose, peptides, and lntralipid. c, Left, heatmap of z-scored activity during IG Ensure infusion in fasted mice. Right, pie charts showing proportions of activated, inhibited, and non-responsive neurons during and after infusion. d, CDF of mean z-scores (0-30 min) across the entire trial during IG infusion of Ensure for fasted versus ad libitum-fed animals. e, CDF of mean z-scores (0-30 min) during IG infusion of glucose, peptides, and lntralipid. f, CDF of time to half-maximal activation for IG infusion of glucose, peptides, and lntralipid. g, Mean z-scores of activated neurons for the entire trial, infusion period (0-10 min), and post-infusion period (10-30 min) for saline, Ensure, glucose, peptides, and lntralipid. h, Percentage of activated neurons across the same three trial epochs. i, Left, Heatmap and PSTH of cNTS neuron activity during IG infusion of methylcellulose. Right, pie charts showing percentage of activated cells during and after IG infusion. j, Heatmap of cNTS responses to saline injection. k, Left, heatmap of responses to Ex-4 injection. Right, pie charts showing percentage of activated cells in the first 10 min and last 20 min. I, Left, heatmap of responses to CCK injection. Right, pie charts showing percentage of activated cells in the first 10 min and last 20 min. m, CDF of mean z-scores (0-30 min) for saline, Ex-4, and CCK injections. n, Heatmaps comparing cNTS neuron activity during IG infusion methylcellulose and subsequent CCK injection. o, Individual neuron traces showing responses to IG infusion of methylcellulose followed by CCK injection. p, Venn diagram showing percentage of cells responsive to methylcellulose, CCK, or both. q, Heatmap of cNTS neuron activity during IG infusion of lntralipid after vehicle injection r, Percentage of activated, inhibited, and non-responsive neurons during and after IG infusion of lntralipid after vehicle or devazepide injection. NS, *P<0.05, **P<0.01, ***P<0.001, ****P<0.0001. Data are mean± sem.

**Extended Data Figure 2:**
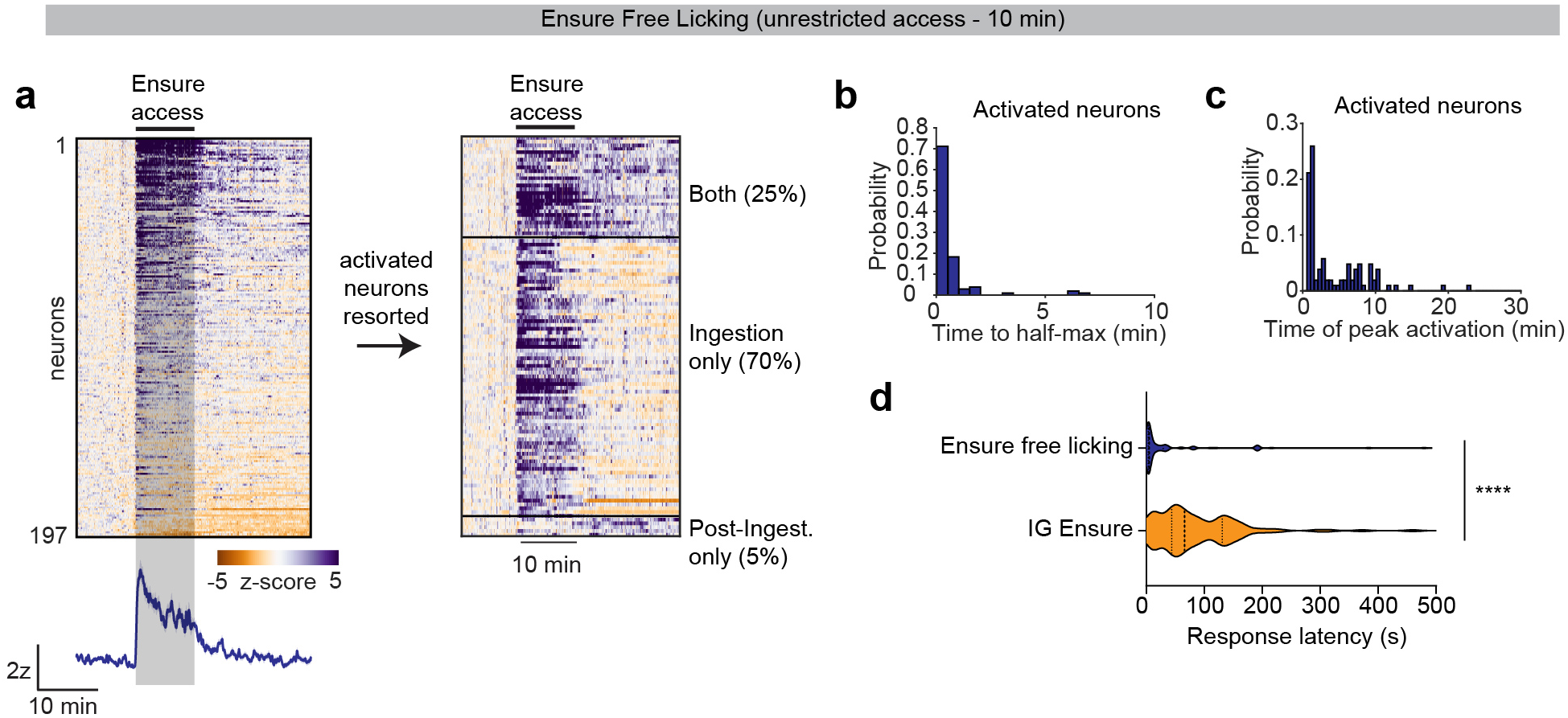
Most cNTS neurons are rapidly activated by self-paced food ingestion. a, Heatmap of cNTS neuron activity during 10 min of self-paced Ensure ingestion. Neurons are sorted by response category (ingestion-activated, post-ingestion-activated, or non-responsive). b, Histogram of time to half-maximal activation for all ingestion-responsive neurons. c, Histogram of time to peak activation during the entire trial (0-30 min). d, Response latency of cNTS neurons during Ensure free licking or IG infusion of Ensure (1 ml). The response latency was calculated using a derivative-based method (See Methods). NS, *P<0.05, **P<0.01, ***P<0.001, ****P<0.0001.

**Extended Data Figure 3:**
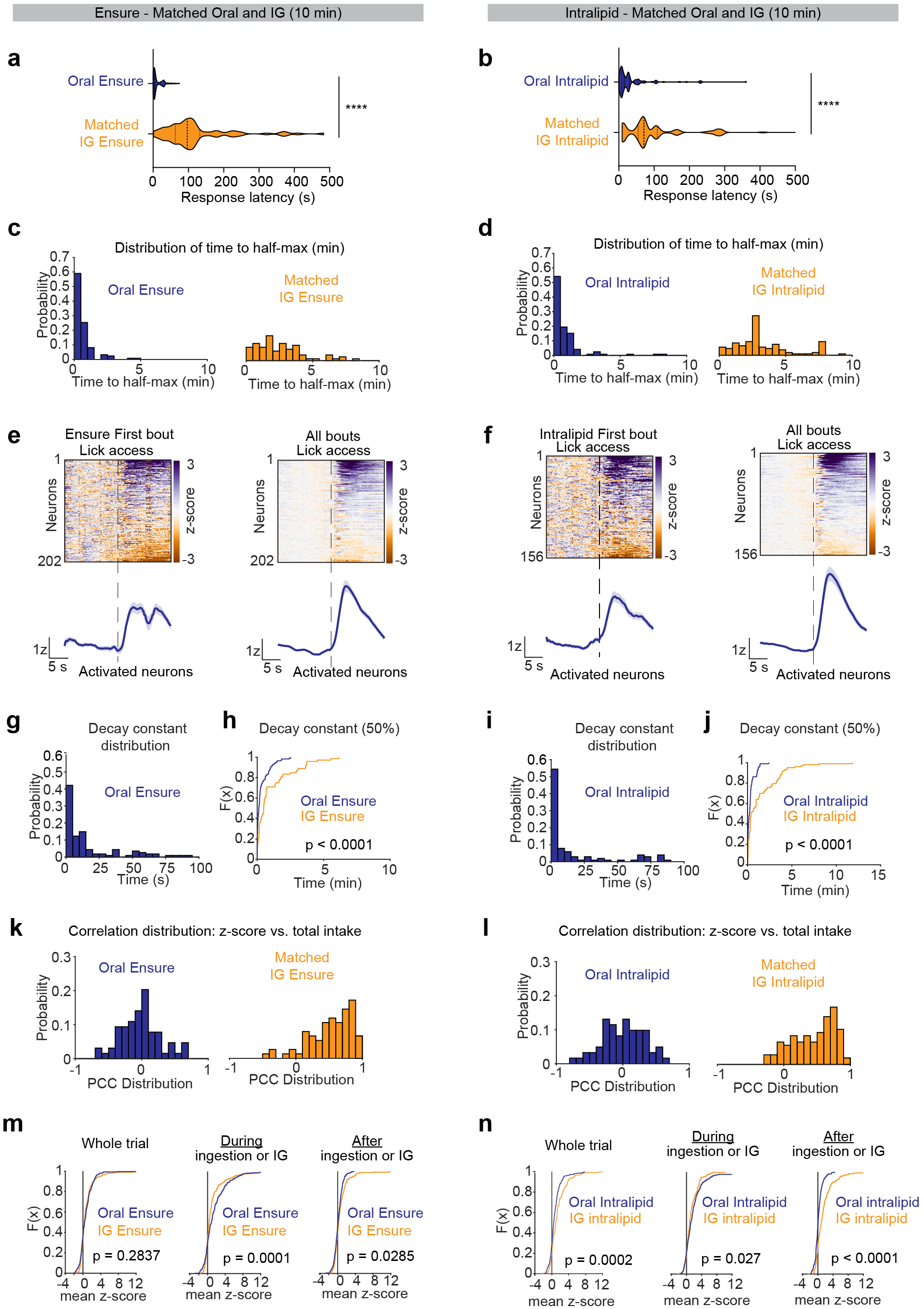
cNTS neuron responses depend on the route of food delivery. a, Response latency of cNTS neurons during oral Ensure ingestion and matched IG infusion of Ensure. The response latency was calculated using a derivative-based method (See Methods). b, Response latency of cNTS neurons during oral lntralipid ingestion and matched IG infusion of lntralipid. c, Histogram of time to half-maximal activation for oral Ensure and matched IG infusion of Ensure. d, Histogram of time to half-maximal activation for oral lntralipid and matched IG infusion of lntralipid. e, Heatmaps and PSTH aligned to first lick for cNTS neurons activated during first-bout lick access and across all bouts for oral Ensure. f, Heatmaps and PSTH aligned to first lick for cNTS neurons activated during first-bout lick access and across all bouts for oral lntralipid. g, Histogram of decay constants (time to 50% of z-score) at the end of oral Ensure ingestion. h, CDF of decay constants for oral versus IG Ensure. i, Histogram of decay constants (time to 50% of z-score) at the end of oral lntralipid ingestion. j, CDF of decay constants for oral versus IG lntralipid. k, Histogram of PCC between z-scored change in activity vs. total intake for oral Ensure and matched IG infusion of Ensure. I, Histogram of PCC between z-scored change in activity vs. total intake for oral lntralipid and matched IG infusion of lntralipid. m, CDF of mean z-scores during the whole trial, ingestion/infusion period, and post-ingestion period for oral Ensure versus matched-volume IG infusion of Ensure. n, CDF of mean z-scores during the whole trial, ingestion/infusion period, and post-ingestion period for oral lntralipid versus matched-volume IG infusion of lntralipid. NS, *P<0.05, **P<0.01, ***P<0.001, ****P<0.0001.

**Extended Data Figure 4:**
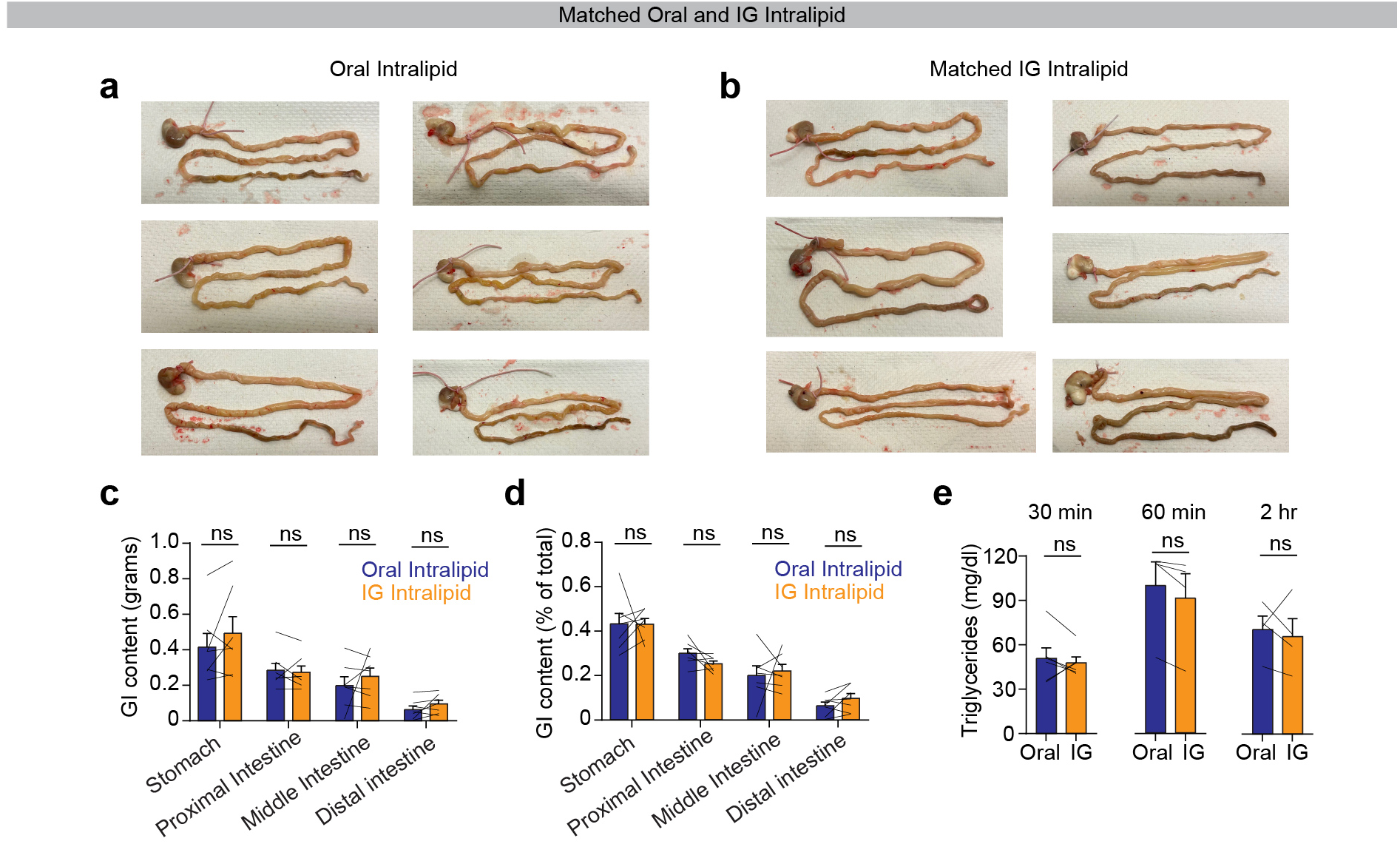
Oral and IG intralipid differences are not due to gross changes in GI motility. a, GI contents after oral lntralipid ingestion. b, GI contents after matched **IG** infusion of lntralipid. c, Quantification of GI content (grams) across stomach and intestinal segments following oral versus matched **IG** infusion of lntralipid d, Percentage of total GI content across stomach and intestinal segments following oral versus matched **IG** infusion of lntralipid. e, Plasma triglyceride concentrations collected at 30, 60, and 120 min following oral versus matched IG infusion of lntralipid. NS, *P<0.05, **P<0.01, ***P<0.001, ****P<0.0001. Data are mean± sem.

**Extended Data Figure 5:**
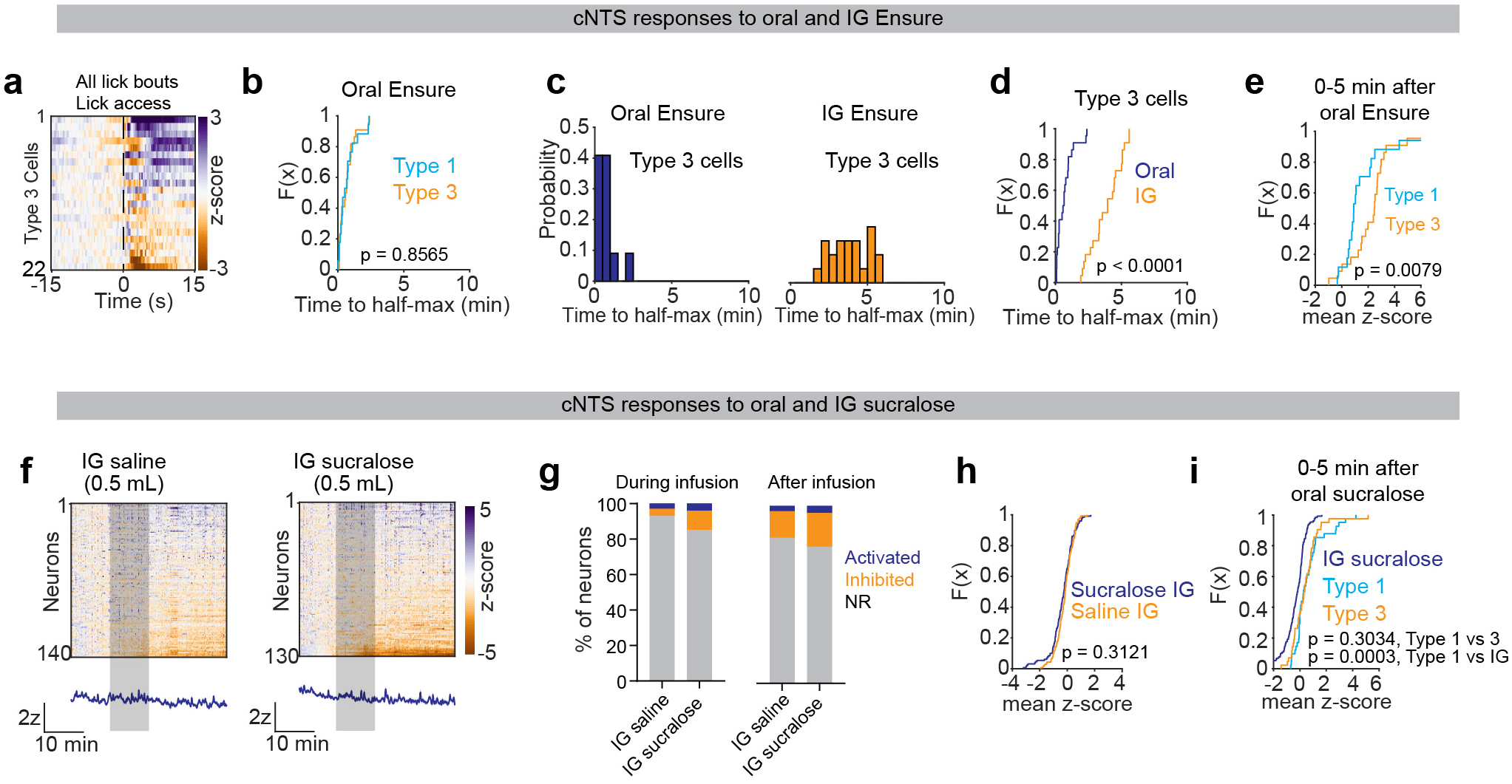
Oral and GI feedback activate partially overlapping ensembles. a, Heatmap of cNTS neuron activity for Type 3 neurons aligned to lick access across all bouts. b, CDF of time to half-max activation for oral Ensure ingestion in Type 1 and 3 neurons. c, Histogram of time to half-max activation for Type 3 neurons during oral Ensure ingestion (left) and IG infusion of Ensure (right). d, CDF of time to half-max activation for oral versus IG Ensure in Type 3 neurons. e, CDF of mean z-scores (0-5 min after oral Ensure ingestion) in Type 1 and 3 neurons. f, Heatmaps of cNTS neuron responses during IG infusion (0.5 ml) of saline (left) and IG infusion of sucralose (right). g, Percentage of neurons classified as activated, inhibited, or non-responsive during IG infusion of saline and sucralose (0-10 min) and after infusion (10-30 min). h, CDF of mean z-scores (0-30 min) during IG infusion of saline and sucralose. i, CDF of mean z-scores (0-5 min) after IG infusion of sucralose or oral sucralose ingestion for Type 1 and 3 neurons. NS, *P<0.05, **P<0.01, ***P<0.001, ****P<0.0001. Data are mean± sem.

**Extended Data Figure 6:**
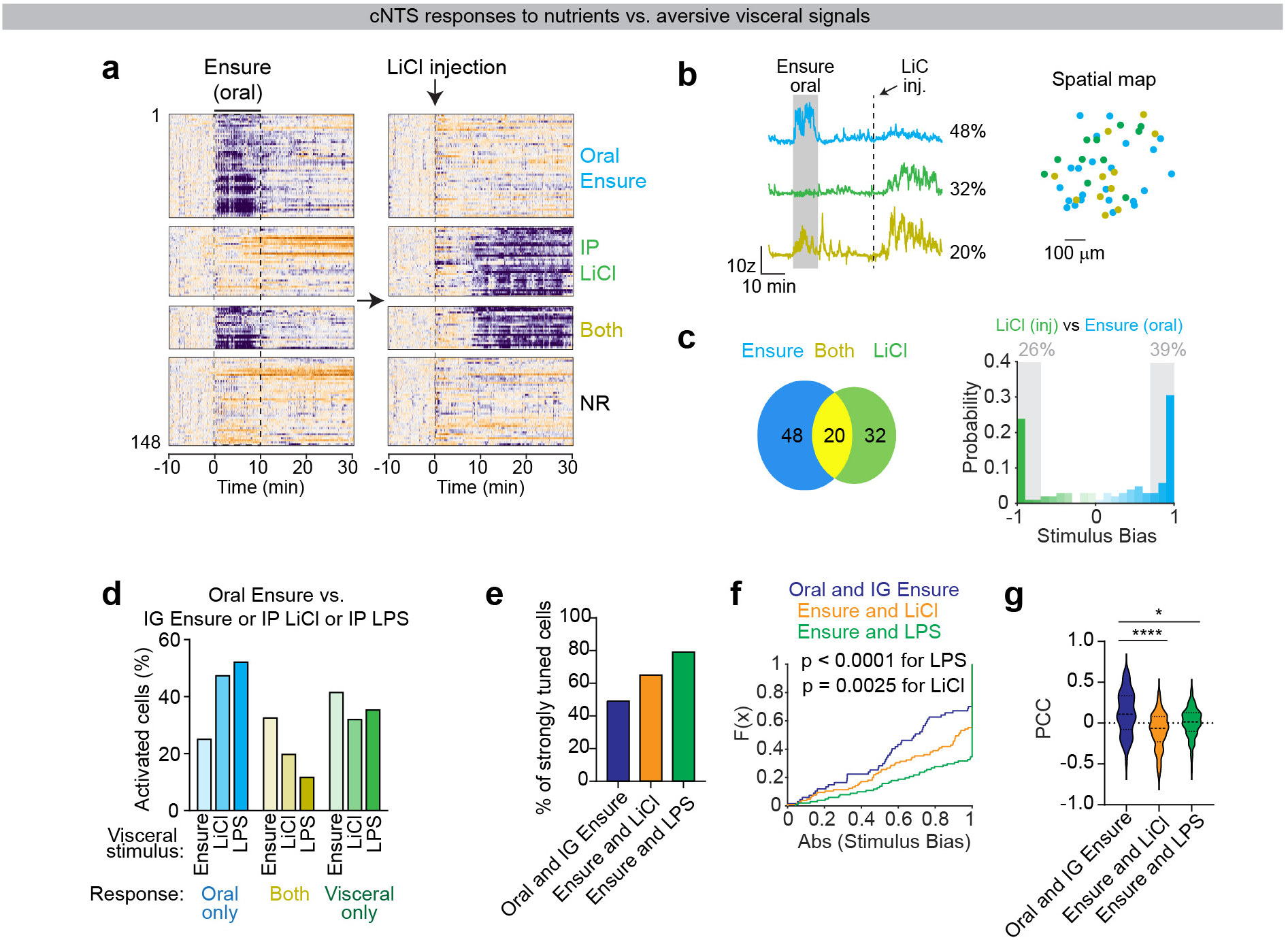
cNTS neurons show sharper tuning between food ingestion and aversive visceral signals. a, Heatmaps of cNTS neuron activity during oral Ensure ingestion (left) and LiCI injection (right). Neurons are categorized as oral-responsive, LiCl-responsive, responsive to both, or non-responsive (NR). b, Left, individual neuron traces for oral Ensure vs. LiCI injection. Right, spatial map of activated neurons for oral Ensure vs. LiCI injection. c, Left, Venn diagram showing percentage of cells activated oral Ensure vs. LiCI injection. Right, histogram of stimulus bias indexes for oral Ensure vs. LiCI injection. d, Percentages of neurons that respond selectively to oral Ensure versus IG infusion of Ensure, LiCI injection, or LPS injection. e, Percentages of neurons classified as strongly tuned (stimulus bias index less than −0.7 or greater than 0.7) for each oral-GI pairing (oral Ensure vs. IG Ensure, oral Ensure vs. LiCI injection, and oral Ensure vs. LPS injection). f, CDF of the absolute value of stimulus bias indexes for oral Ensure against IG Ensure, LiCI injection, or LPS injection. cNTS neurons show sharpest tuning between oral Ensure and LPS injection. g, Pearson correlation coefficient (PCC) for the z-scored change in activity between oral Ensure versus IG Ensure, LiCI injection, or LPS injection. NS, *P<0.05, **P<0.01, ***P<0.001, ****P<0.0001. Data are mean± sem.

**Extended Data Figure 7:**
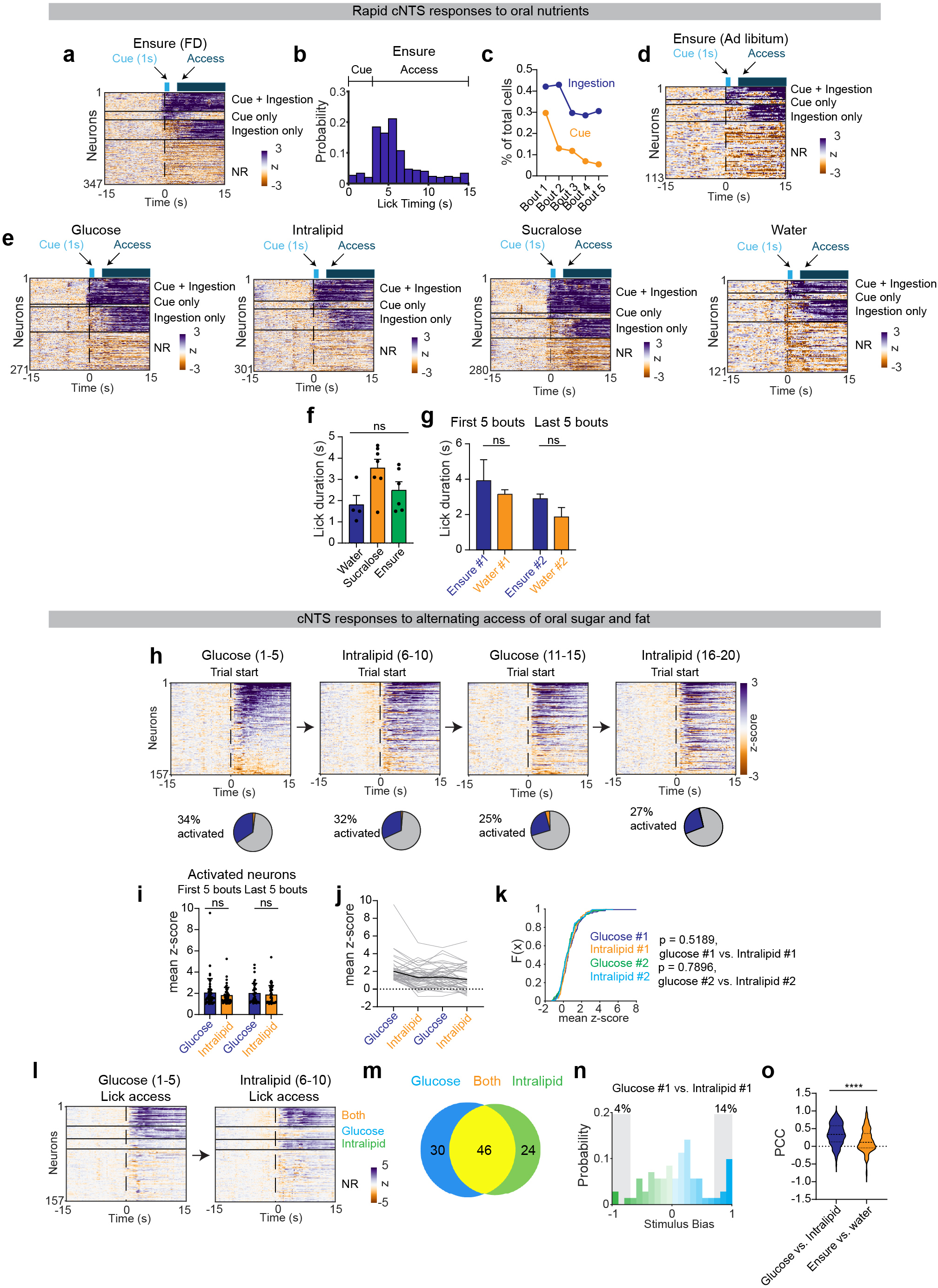
Regulation of cNTS neurons by oral nutrients and food-associated tastes. a, Heatmap of cNTS neuron activity aligned to trial start during oral Ensure ingestion in fasted mice, with neurons categorized as responsive to cue+ingestion, cue-only, ingestion-only, or non-responsive. b, Histogram of when licks occur aligned to trial start for Ensure ingestion. c, Percentage of activated cells during the cue or ingestion segment for the first 5 trials. d, Heatmap of cNTS neuron activity aligned to trial start during oral Ensure ingestion in ad-libitum fed mice, with neurons categorized as responsive to cue+ingestion, cue-only, ingestion-only, or non-responsive. e, Heatmap of cNTS neuron activity aligned to trial start during oral ingestion of glucose, lntralipid, sucralose, or water in fasted mice, with neurons categorized as responsive to cue+ingestion, cue-only, ingestion-only, or non-responsive. f, Lick duration for water, sucralose, and Ensure ingestion across the first five bouts. g, Lick duration between the first five and last five bouts for Ensure and water ingestion. h, Heatmaps of cNTS neuron activity during alternating trials of oral glucose (trials 1-5), lntralipid (6-10), glucose (11-15), and lntralipid (16-20). Pie charts indicate percentage of activated neurons per block. i, Mean z-scores of activated neurons during the first five and last five bouts across alternating glucose and lntralipid trials. j, Mean z-score responses of individual neurons tracked across blocks of glucose or lntralipid consumption k, CDF of mean z-scores (0-15s) for glucose vs. lntralipid across the first and last blocks. I, Heatmaps of lick-aligned responses to glucose (trials 1-5) and lntralipid (6-10), with neurons grouped as responsive to glucose only, lntralipid only, both, or non-responsive. m, Venn diagram showing percentage of cells activated by oral glucose vs. oral lntralipid. n, Histogram of stimulus bias indexes for the first block of oral glucose and lntralipid ingestion. o, Pearson correlation coefficients **(PCC)** for z-scored change in activity between glucose and lntralipid ingestion, or Ensure and water ingestion. The dynamics of glucose and lntralipid ingestion are more correlated. NS, *P<0.05, **P<0.01, ***P<0.001, ****P<0.0001. Data are mean± sem.

**Extended Data Figure 8:**
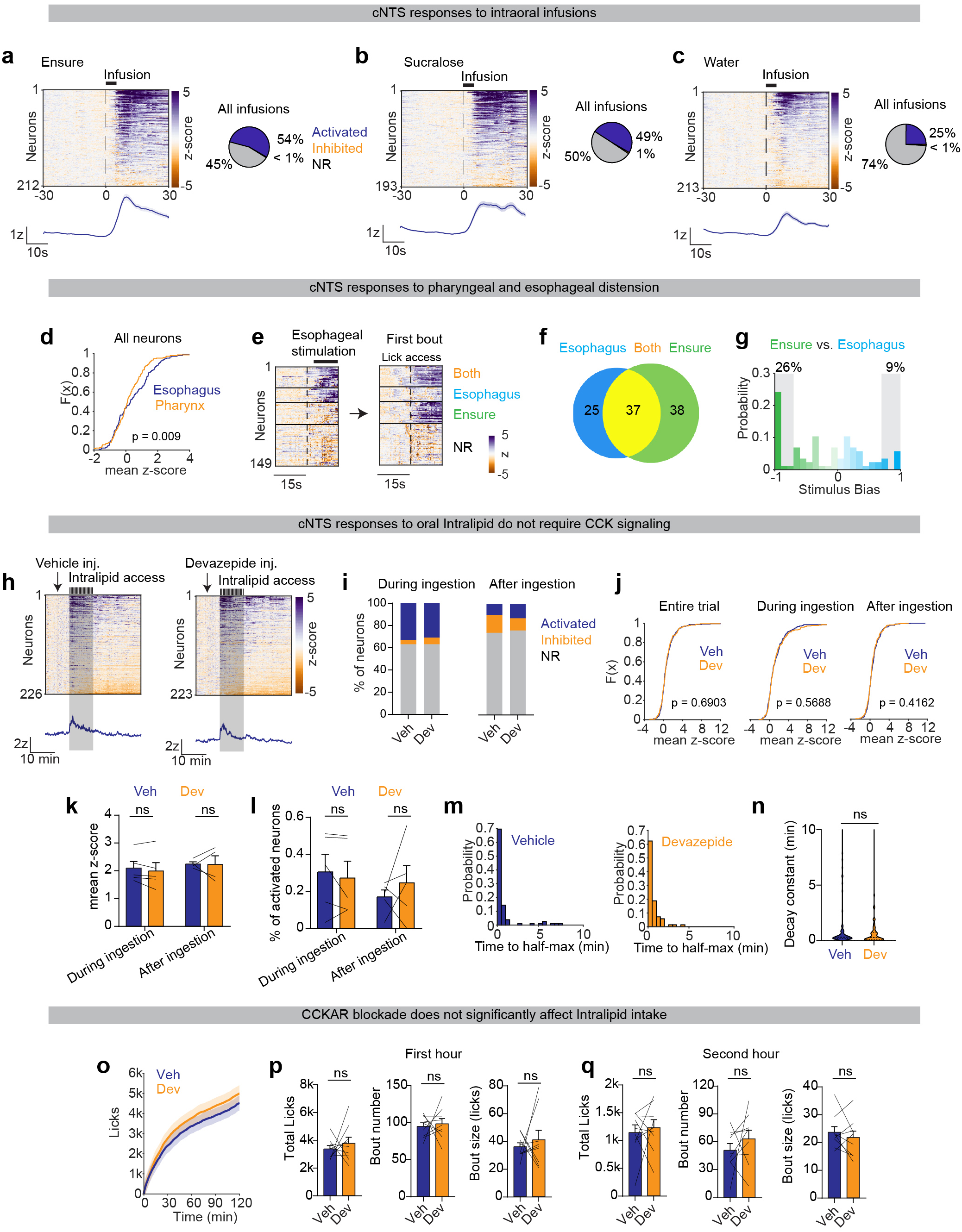
Orosensory mechanisms for rapid cNTS activation during food ingestion. a, Left, Heatmaps of cNTS neuronal responses during intraoral infusions of Ensure. PSTH aligned to infusion start below heatmap. Right, pie chart show percentage of activated, inhibited, and non-responsive neurons across all intraoral trials. b, Left, Heatmaps of cNTS neuronal responses during intraoral infusions of sucralose. PSTH aligned to infusion start below heatmap. Right, pie chart show percentage of activated, inhibited, and non-responsive neurons across all intraoral trials. c, Left, Heatmaps of cNTS neuronal responses during intraoral infusions of water. PSTH aligned to infusion start below heatmap. Right, pie chart show percentage of activated, inhibited, and non-responsive neurons across all intraoral trials. d, CDF of mean z-scores (0-15s) for cNTS neuron responses to pharyngeal versus esophageal distension. e, Heatmaps showing neuronal responses to esophageal distension (left) and to the first oral Ensure bout (right) in the same animal. f, Venn diagram showing percentage of neurons responsive to esophageal distension, oral Ensure, or both. g, Histogram of stimulus bias indexes for esophageal distension vs. the first oral Ensure trial. h, Heatmaps of neuronal responses during oral lntralipid ingestion following vehicle injection (left) or devazepide injection (right). PSTH aligned to lntralipid access are below each heatmap. i, Percentage of activated, inhibited, and non-responsive neurons during oral lntralipid ingestion (left) and post-ingestion (right) after vehicle or devazepide injection. j, CDF of mean z-scores across the entire trial, during ingestion (0-10 min), and after ingestion (10-30 min) for vehicle or devazepide injection. k, Mean z-scores of activated neurons during oral lntralipid ingestion and post-ingestion after vehicle or devazepide injection. I, Percentage of activated neurons in the ingestion and post-ingestion periods after vehicle or devazepide injection. m, Histogram of time to half-maximal activation during oral lntralipid ingestion after vehicle (left) or devazepide injection (right). n, Time to half-decay during oral lntralipid ingestion after vehicle or devazepide injection. o, Cumulative licks during 2-hour oral lntralipid access following vehicle or devazepide injection. p, Total licks, bout number, and bout size (licks) during the first hour of lntralipid ingestion after vehicle or devazepide injection. q, Total licks, bout number, and bout size (licks) during the second hour of lntralipid ingestion after vehicle or devazepide injection. NS, *P<0.05, **P<0.01, ***P<0.001, ****P<0.0001. Data are mean± sem.

**Extended Data Figure 9:**
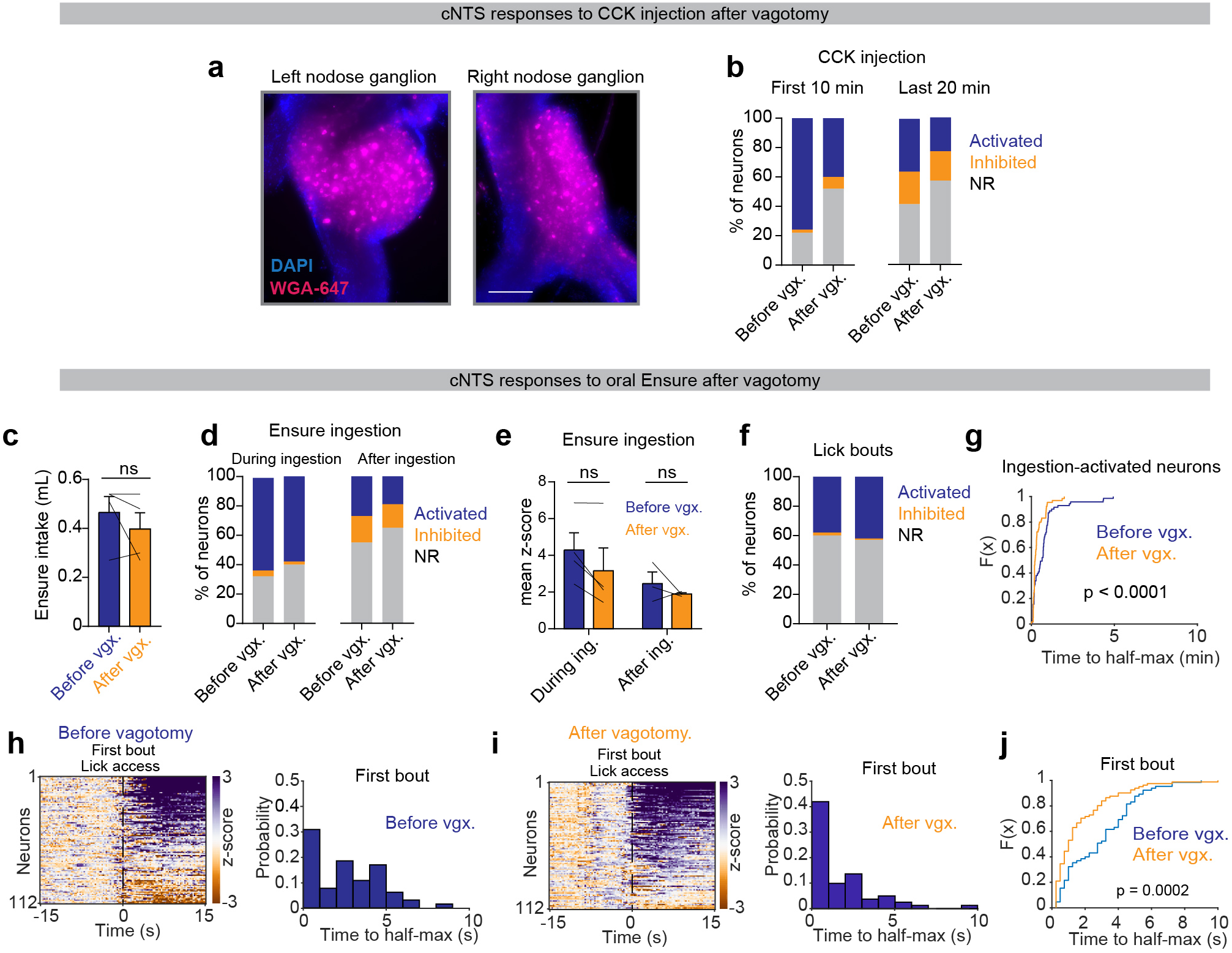
The vagus nerve is dispensable for rapid cNTS activation by food ingestion. a, Histological validation of left and right nodose ganglia labeled by WGA-647 (magenta) and DAPI (blue) in a control Vglut2Cre animal without vagotomy. b, Percentage of activated, inhibited, and non-responsive neurons during the first 10 min (left) and last 20 min (right) following CCK injection before and after vagotomy. c, Oral Ensure intake in the same mice before and after vagotomy. d, Percentage of neurons classified as activated, inhibited, or non-responsive during oral Ensure ingestion (0-10 min) or after ingestion (10-30 min), before and after vagotomy. e, Mean z-scores of activated neurons during ingestion and post-ingestion periods before and after vagotomy. f, Percentage of activated, inhibited, and non-responsive neurons during lick bouts (0-15s) before versus after vagotomy. g, CDF of time to half-maximal activation for ingestion-activated neurons before and after vagotomy. h, Heatmap of cNTS neuron activity during the first Ensure lick bout before vagotomy (left) and corresponding histogram of time to half-max (right). i, Heatmap of cNTS neuron activity during the first Ensure lick bout after vagotomy (left) and corresponding histogram of time to half-max (right). j, CDF of time to half-max activation for first-bout Ensure responses before versus after vagotomy. NS, *P<0.05, **P<0.01, ***P<0.001, ****P<0.0001. Data are mean± sem.

**Extended Data Figure 10:**
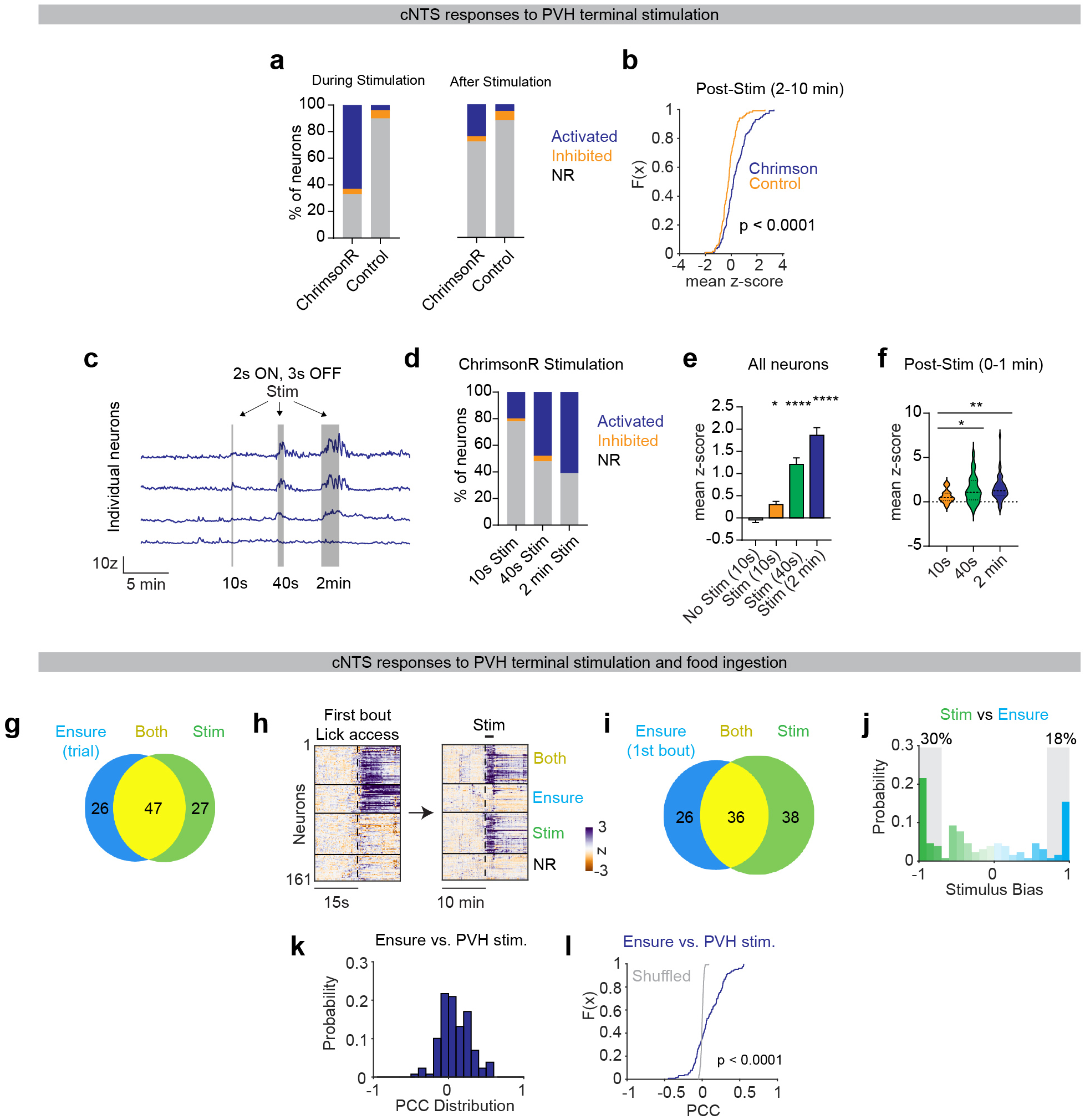
PVH terminals activate downstream cNTS neurons. a, Percentages of cNTS neurons classified as activated, inhibited, or non-responsive during PVH + cNTS ChrimsonR photostimulation (left) and in the post-stimulation period (right), compared to no-opsin controls. b, CDF of mean z-scores during the 2-10 min post-stimulation window for ChrimsonR versus control mice. c, Individual cNTS neuron traces during 2s ON/ 3s OFF photostimulation for stimulation durations of 1Os, 40s, and 2 min. d, Percentage of neurons categorized as activated, inhibited, or non-responsive at each stimulation duration in ChrimsonR mice. e, Mean z-scores across all neurons during stimulation for no-stim controls and each stimulation duration. f, Mean z-scores in the immediate post-stimulation period (0-1 min) for each stimulation duration. g, Venn diagram percentage of cNTS neurons responsive to oral Ensure ingestion (0-10 min), PVH terminal stimulation, or both. h, Heatmaps of cNTS neuron activity aligned to the first Ensure lick bout (left) and to PVH terminal stimulation (right), with neurons categorized as responsive to Ensure only, stimulation only, both, or non-responsive. i, Venn diagram showing percentage of neurons responsive to the first Ensure bout (0-15s), PVH terminal stimulation, or both. j, Histogram of stimulus bias indexes for the first bout of Ensure ingestion vs. PVH terminal stimulation. k, Histogram of Pearson correlation coefficients (PCC) between single-cell responses to Ensure ingestion and PVH terminal stimulation. I, CDF of PCC for Ensure ingestion versus PVH terminal stimulation with shuffled controls (gray). NS, *P<0.05, **P<0.01, ***P<0.001, ****P<0.0001. Data are mean± sem.

**Extended Data Figure 11:**
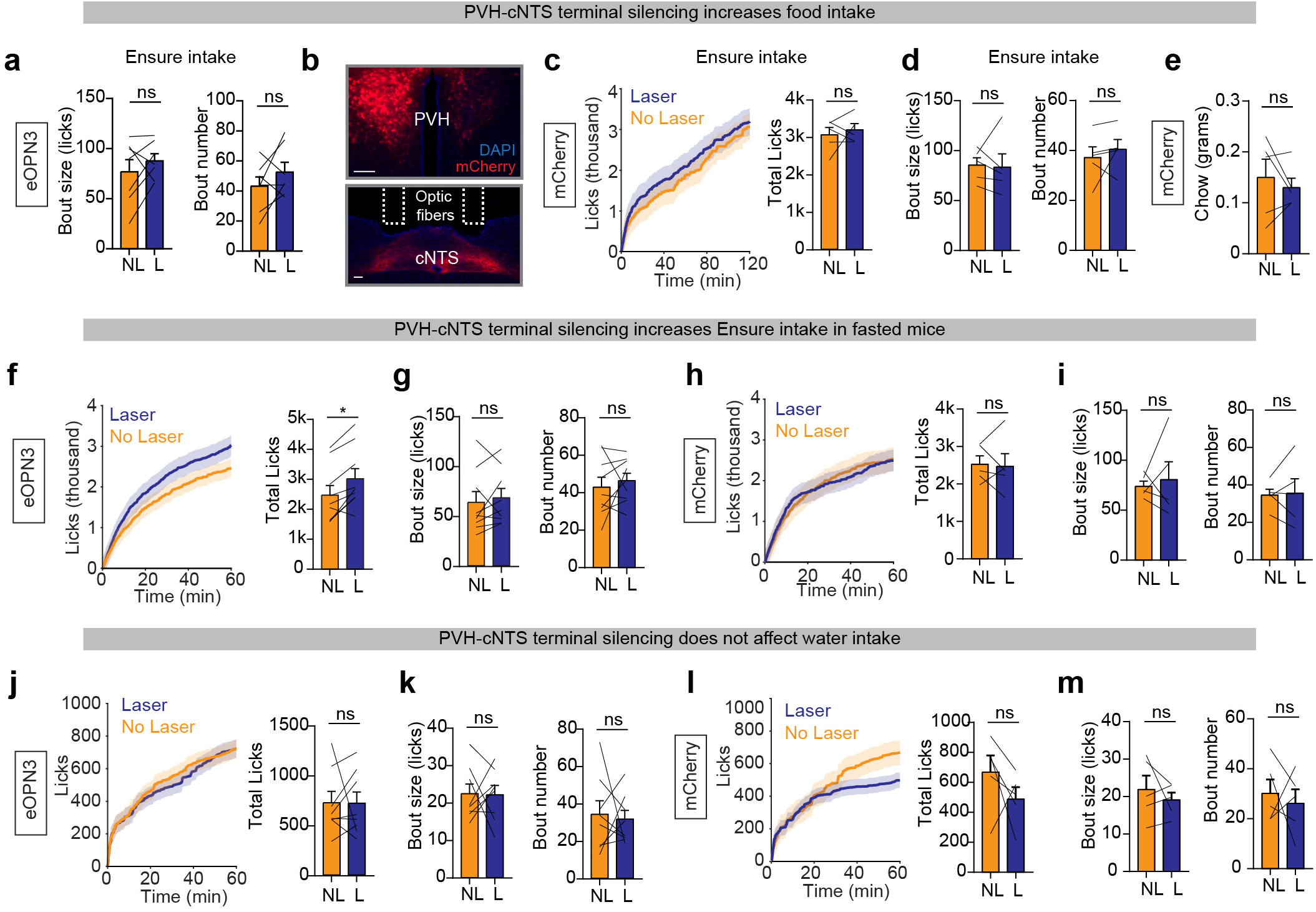
PVH---+cNTS terminal silencing increases food intake. a, Bout size (left) and bout number (right) during dark-phase Ensure intake (2 hours) for mice expressing eOPN3 with no laser (NL) versus laser (L). b, Histological verification of PVH neurons and terminals in the cNTS expressing mCherry and optic fiber placement above the cNTS. Scale bar= 100 µm. c, Left, cumulative licks during dark-phase Ensure intake for NL versus L. Right, total licks. d, Bout size (left) and bout number (right) during dark-phase Ensure intake (2 hours) for mice expressing mCherry with NL versus L. e, Total chow consumed (grams) during the dark phase for mice expressing mCherry for NL versus L. f, Left, cumulative licks during Ensure intake for fasted mice expressing eOPN3 with NL versus L. Right, total licks. g, Bout size (left) and bout number (right) during Ensure intake (1 hour) for fasted mice expressing eOPN3 with NL versus L. h, Left, cumulative licks during Ensure intake for fasted mice expressing mCherry with NL versus L. Right, total licks. i, Bout size (left) and bout number (right) during Ensure intake (1 hour) for fasted mice expressing mCherry with NL versus L. j, Left, cumulative licks during water intake for water-deprived mice expressing eOPN3 with NL versus L. Right, total licks. k, Bout size (left) and bout number (right) during water intake (1 hour) for water-deprived mice expressing eOPN3 with NL versus L. I, Left, cumulative licks during water intake for water-deprived mice expressing mCherry with NL versus L. Right, total licks. m, Bout size (left) and bout number (right) during water intake (1 hour) for water-deprived mice expressing mCherry with NL versus L. NS, *P<0.05, **P<0.01, ***P<0.001, ****P<0.0001. Data are mean± sem.

